# Trabecular bone remodeling in the ageing mouse: a micro-multiphysics agent-based *in silico* model using single-cell mechanomics

**DOI:** 10.1101/2022.11.16.516723

**Authors:** Daniele Boaretti, Francisco C. Marques, Charles Ledoux, Amit Singh, Jack J. Kendall, Esther Wehrle, Gisela A. Kuhn, Yogesh D. Bansod, Friederike A. Schulte, Ralph Müller

## Abstract

Bone remodeling is regulated by the interaction between different cells and tissues across many spatial and temporal scales. Notably, *in silico* models are regarded as powerful tools to further understand the signaling pathways that regulate this intricate spatial cellular interplay. To this end, we have established a 3D multiscale micro-multiphysics agent-based (micro-MPA) *in silico* model of trabecular bone remodeling using longitudinal *in vivo* data from the sixth caudal vertebra (CV6) of PolgA^(D257A/D257A)^ mice, a mouse model of premature aging. Our model includes a variety of cells as single agents and receptor-ligand kinetics, mechanomics, diffusion and decay of cytokines which regulate the cells’ behavior. We highlighted its capabilities by simulating trabecular bone remodeling in the CV6 of 5 mice over 4 weeks and we evaluated the static and dynamic morphometry of the trabecular bone microarchitecture. Based on the progression of the average trabecular bone volume fraction (BV/TV), we identified a configuration of the model parameters to simulate homeostatic trabecular bone remodeling, here named basal. Crucially, we also produced anabolic, anti-anabolic, catabolic and anti-catabolic responses with an increase or decrease by one standard deviation in the levels of osteoprotegerin (OPG), receptor activator of nuclear factor kB ligand (RANKL), and sclerostin (Scl) produced by the osteocytes. Our results showed that changes in the levels of OPG and RANKL were positively and negatively correlated with the BV/TV values after 4 weeks in comparison to basal levels, respectively. Conversely, changes in Scl levels produced small fluctuations in BV/TV in comparison to the basal state. From these results, Scl was deemed to be the main driver of equilibrium while RANKL and OPG were shown to be involved in changes in bone volume fraction with potential relevance for age-related bone features. Ultimately, this micro-MPA model provides valuable insights into how cells respond to their local mechanical environment and can help to identify critical pathways affected by degenerative conditions and ageing.

## Introduction

With aging, bone becomes more fragile with increasing fracture risk, leading to an increased morbidity and mortality (Kanis et al. 2021). To better understand how aging affects bone cellular behavior, *in vivo* and *in silico* studies have been used to analyze the tissue and cellular properties of bone mechanobiology. It is known that bone remodeling is regulated by the interaction between different cells and tissues (i.e. cancellous, cortical and marrow tissues) across many spatial and temporal scales. Hybrid modeling has been identified as a powerful modeling technique which combines multiscale, multiphysics, and agent-based modeling to analyze in a unified way the organ, tissue, cell, and gene scales over weeks, months or even years (Boaretti et al. 2022). Such an *in silico* model can deal with the action of single cells while millions of cells are active and interact with each other in a multiscale simulation, making this tool ideal to explore the hierarchical processes governing bone mechanobiology. Furthermore, by enabling fine modelling of signaling pathways that regulate the cells’ activity, such models can help to estimate physiological cytokine production rates and explore their effect on spatial cellular distribution, bone formation, and bone resorption, which is hardly feasible *in vivo*. Yet, bone remodeling has not been quantitatively studied *in silico* across spatial and temporal scales and using 3D *in vivo* data as reference.

Recent experimental studies have analyzed the effects of aging in mice and how mechanical loading can counterbalance frailty (Scheuren et al. 2020a, b). Accelerated aging was provoked in PolgA^(D257A/D257A)^ mice due to systemic mitochondrial dysfunction caused by accumulation of mitochondrial DNA point mutations (Trifunovic et al. 2004; Kujoth et al. 2005). The findings from these publications have mainly focused on cortical and trabecular bone morphometric parameters with analyses of selected gene and protein expression on specific time points. The study of Dobson and colleagues (Dobson et al. 2020) showed that in this PolgA^(D257A/D257A)^ mouse model, osteoblast density was reduced and the production of mineralized matrix by osteoblasts was significantly impaired, leading to reduced bone formation rates. The mechanisms which lead to cellular alterations in bone with aging remain to be further elucidated.

We have recently proposed an *in silico* multiscale micro-multiphysics agent-based (micro-MPA) model (Boaretti et al. 2018) adapted from the model of fracture healing in cortical bone by Tourolle (2019). This micro-MPA model was based on multiscale (from the organ to the protein spatial scales and from weeks to minutes as temporal scales), multiphysics (mechanical signal, reaction-diffusion-decay of cytokines), and agent-based modeling (single-cell behavior). Each cell was represented as a single agent, and signaling pathways were modeled to regulate the cell behavior with distinct expression and bone remodeling profiles dependent on the mechanical signal which we define as mechanomics. Crucially, this enabled simulating the movement, differentiation, cluster behavior, apoptosis of bone cells, their response to the local mechanical environment in terms of synthesis of cytokines and bone remodeling in the trabecular region, employing the same isotropic voxel resolution of the 3D micro-computed tomography (micro-CT) *in vivo* data. In this model, osteoprotegerin (OPG), receptor activator of nuclear factor kB ligand (RANKL), sclerostin (Scl) were cytokines that decayed, diffused in the volume, and were produced by the cells. Transforming growth factor beta 1 (TGF-β1) was modeled as cytokine stored in the bone volume and it could diffuse and decay after resorption. The signaling pathways were modeled with receptor-ligand kinetics, with the receptors located on the cells’ surfaces. In addition, a free ligand could bind to another free ligand, e.g., OPG can bind to RANKL. These pathways have been shown to be the main regulators of cell differentiation and proliferation (Krishnan et al. 2006; Boyce and Xing 2008; Lin et al. 2009; Tang et al. 2009; Warren et al. 2015; Elson et al. 2022). The osteocytes were considered the main mechanosensors of the mechanical signal (Santos et al. 2009; Klein-Nulend et al. 2012, 2013) and they promoted bone formation or resorption by releasing cytokines into the volume to regulate the signaling pathways affecting eventually the osteoblasts and osteoclasts. These cells formed and resorbed bone on the surface, respectively. Osteocytes, osteoblasts and osteoclasts were the basis for the regulation of bone remodeling we could simulate using *in vivo* data. Indeed, a previous version of this model was used for simulating denosumab treatment in human biopsies over 10 years, showing the potential of micro-MPA models for designing optimal clinical trials (Tourolle et al. 2021).

In the current study, we present an updated micro-MPA model to simulate how bone adapts and remodels through single-cell mechanomics in mice. We hypothesized that cells produce cytokines to regulate the cellular actions as a response to the local mechanical signal they perceive, e.g., osteocytes and osteoblasts release cytokines to promote anabolic or anti-catabolic responses under high effective strain (EFF) and osteocytes and osteoblasts release catabolic or anti-anabolic cytokines under low EFF. This hypothesis is implemented with an adaptation of the bone microarchitecture through an EFF-based sigmoidal dose-response mechanomics at the cellular level. Furthermore, we hypothesized that osteocytes are mainly responsible for regulating the other cells’ activity through their single-cell mechanomics.

The aim of this study is to show that the proposed *in silico* model can reproduce a homeostatic condition similar to the longitudinal *in vivo* data, where a dynamic equilibrium between bone formation and bone resorption maintains relatively constant bone volume fraction (Rodan 1998; Nakahama 2010; Rauner et al. 2020). Moreover, we investigated the quantitative effect of manipulating the cytokines levels involved in the signaling pathways on the bone morphometric parameters. For this purpose, the proposed novel micro-MPA *in silico* model of bone remodeling was applied to micro-CT *in vivo* data to test the effect of different production values of RANKL, OPG, and Scl by osteocytes on the static and dynamic bone morphometry data relatively to the homeostatic configuration.

## Materials and Methods

### *In vivo* input data

*In vivo* data of the control group of a study analyzing the effects of frailty and osteosarcopenia on the bone microarchitecture of prematurely aged PolgA^(D257A/D257A)^ mice (n=9) were used (Scheuren et al. 2020a). Briefly, at an age of 35 weeks, stainless steel pins were inserted at the sixth caudal vertebra (CV6). At week 38, a sham (0N control) loading regime was applied three times per week over 4 weeks. Specifically, mice were placed on the loading machine for 5 minutes, without any mechanical loading. The *in vivo* micro-CT images (vivaCT 40, Scanco Medical AG) were acquired and analyzed every week at an isotropic voxel resolution of 10.5 um. The acquired images showed a slight reduction in the normalized trabecular bone volume fraction over the course of the experiment (2% at the end). All mouse experiments reported in the present study were previously carried out in strict accordance with the recommendations and regulations in the Animal Welfare Ordinance (TSchV 455.1) of the Swiss Federal Food Safety and Veterinary Office, and results are reported following the principles of the ARRIVE guidelines (www.nc3rs.org.uk/arrive-guidelines).

### *In silico* micro-multiphysics agent-based model

In the present study, we adapted the micro-MPA model of Tourolle et al. (2021) for the simulation of homeostatic bone remodeling in the mouse vertebra, including the bone response to physiological loading. A detailed mathematical description of the model was provided by Tourolle et al. (2021). The overview of the adapted *in silico* model is shown in Figure 1.

**Figure 1.**
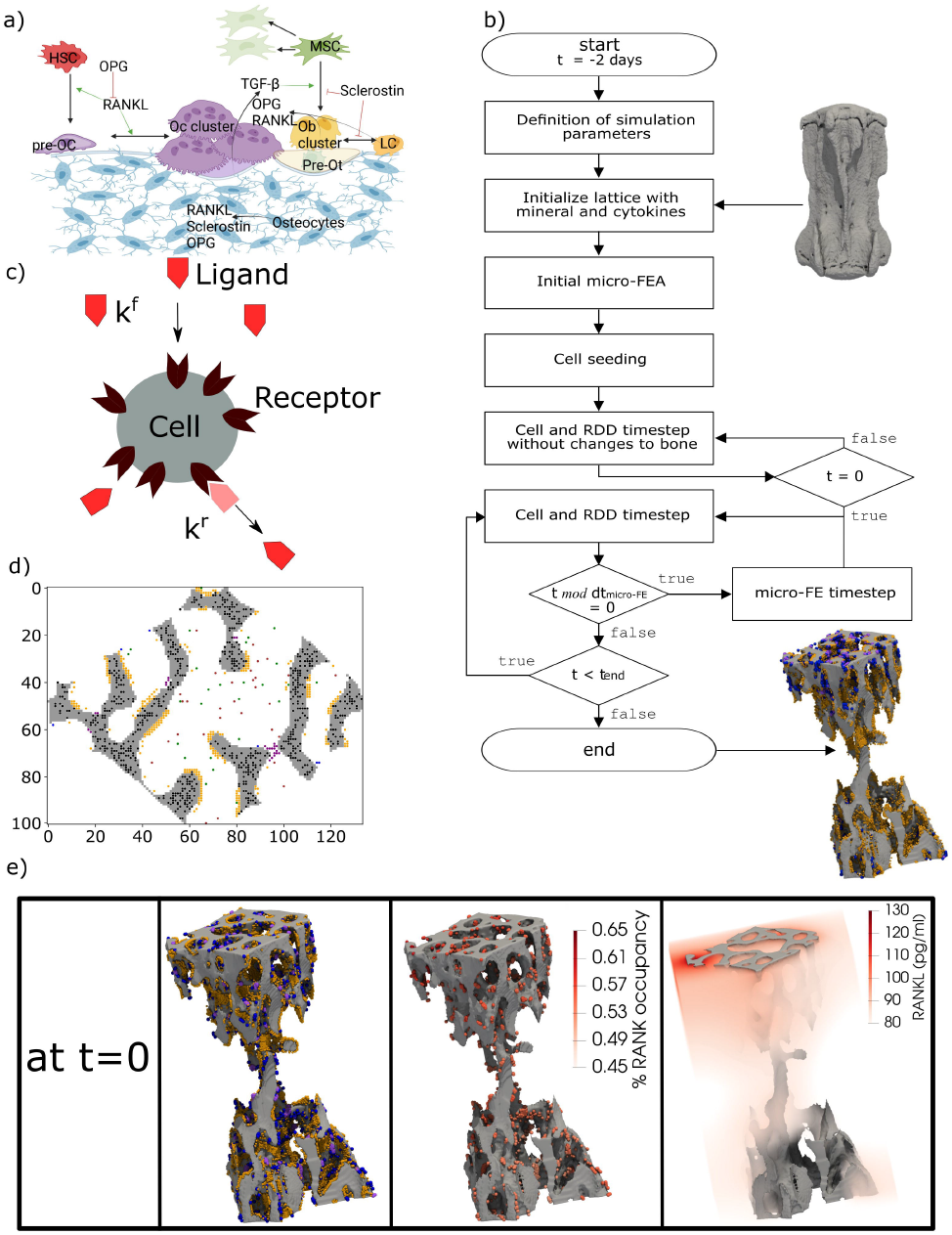
The concept of the micro-MPA *in silico* model in the mouse vertebra. (**A**) The cells and cytokines modeled in the bone remodeling version of the model. (**B**) Overview of the simulation pipeline from the input data to the end of the simulation. (**C**) A schematic representation of the cell receptor-ligand kinetics modelled *in silico*. Here, the ligand can bind only to the targeted cell receptor with a forward rate k^f^ and it can dissociate from the receptor with a backward rate k^r^. This is an example of the modelled receptor-ligand kinetics of Lipoprotein receptor-related protein 5/6 (LRP5/6)-sclerostin for an osteoblast, lining cell and mesenchymal stem cell. (**D**) A 2D slice from the trabecular region of an *in silico* simulation with osteocytes (black), osteoclasts (purple), preosteoclasts (blue), osteoblasts (orange), mesenchymal stem cells (green) and hematopoietic stem cells (brown). (**E**) A snapshot of the initial configuration of the simulation at t=0. On the left, the osteoblasts (orange), osteoclasts (purple) and preosteoclasts (blue) are shown on the trabecular bone surface, in the middle the spatial distribution of the receptor activator of nuclear factor kB (RANK) binding site occupancy on the osteoclasts and preosteoclasts and on the right the spatial distribution of the RANK ligand (RANKL) configuration in the trabecular region. (**A**) created with BioRender.

In the current implementation, the cells are modeled individually using the agent-based paradigm, where all cells of the same type share the same biological properties and follow the same prescribed rules. The model implements the actions of each cell according to the local physiological, chemical, and mechanical environment at the cellular spatial and temporal scale. In our implementation, the changes in the bone microarchitecture are accumulated over the whole time of the experiment (4 weeks) which is much longer than the single temporal step of the cells (40 minutes). The model simulated bone remodeling only within the trabecular region, while the cortical region was kept constant during the simulation, as performed in previous studies (Schulte et al. 2013b; Levchuk et al. 2014; Levchuk 2015).

### Modeling the cellular behavior

Osteocytes (Ot) are embedded in bone and produce RANKL, OPG, and Scl depending on the local mechanical signal they perceive (Santos et al. 2009; Klein-Nulend et al. 2012, 2013). Osteoblasts (Ob) produce osteoid, unmineralized matrix, in their neighborhood towards the surface of the bone, based on the local EFF they perceive. Moreover, Ob may become pre-Ot when the voxel they reside in is at least 50% full of osteoid and there is an osteocyte in the normal direction towards the bone (Franz-Odendaal et al. 2006). Lining cells are considered osteoblasts precursors. In addition, Ob and lining cells also produce OPG and RANKL according to the mechanical signal they perceive. Mesenchymal stem cells (MSC) are present in the marrow space where they can move, proliferate, undergo apoptosis, and can differentiate into an Ob or a lining cell if they are close to the surface and close to an osteoclast (Oc). MSC, lining cells, and Ob have the Lipoprotein receptor-related protein 5/6 (LRP5/6) receptor which binds to Scl (Li et al. 2005; Bourhis et al. 2011). If the bound receptor is higher than a user-defined threshold value (Supplementary Table S1), Ob differentiate into lining cell and MSC differentiate into lining cell if they meet the condition mentioned above. MSC and Ob have also the TGF-β1 receptor on their surface, and if that receptor is highly bound then they can proliferate more frequently. Pre-Ot differentiate into Ot if the voxel they reside in is at least 50% full of mineralized matrix (Franz-Odendaal et al. 2006). The hematopoietic stem cells (HSC) are present in the marrow space, they can move, proliferate, undergo apoptosis and differentiate into pre-Oc if their RANK receptors are highly bound to RANKL (Nelson et al. 2012; Warren et al. 2015). Pre-Oc are motile and can differentiate back into HSCs if the RANK receptor is not highly bound. Moreover, they can differentiate into Oc if there are at least 3 osteoclastic cells (pre-Oc or Oc) in their neighborhood. Oc resorb bone towards the bone surface, with the direction defined by the gradient of the mineral concentration. We calibrated the original implementation of the mineralization kinetics by Tourolle (2019) to have more stable formation rates for simulations of bone remodeling over 4 weeks. We modeled the mineralization kinetics of the matrix in each voxel by changing its mineral concentration to reach its osteoid concentration. The mineralization of a single voxel follows this equation:

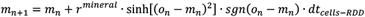

where *m*_*n*_ is the mineral concentration in the voxel at the iteration n, *o*_*n*_ is the osteoid concentration in the voxel at the iteration n, *r*^*mineral*^ is the mineralization rate, *sgn* is the sign function and *dt*_*cells*−*RDD*_ is the timestep for modeling the cellular behavior and the reaction-diffusion-decay step (RDD). With this mineralization rule, the mineral concentration can increase over the iterations if there is a higher concentration of osteoid than the mineral concentration in that voxel, therefore the upper limit of the mineral concentration with this mineralization rule is the osteoid concentration at that voxel. However, the osteoid can increase if there are any osteoblasts in that neighborhood that can release osteoid around themselves. Therefore, given a voxel, if there is more osteoid than mineral, the mineral concentration will increase and if there is more mineral than osteoid, the mineral concentration will decrease. In this way, when Ob release osteoid, there is a delay in bone formation because osteoid leads to a change in the mineral concentration through the mineralization process. In this model, if pre-Ot becomes Ot because the mineral concentration is at least 0.5, that voxel where the cell is present is considered bone, but its mineral concentration does not jump to 100%.

To simulate trabecular bone remodeling, we modeled TGF-β1, RANKL-RANK-OPG, and LRP5/6-Scl signaling pathways at the receptor-ligand level and the Ob, lining cell, Oc, pre-Oc, Ot, pre-Ot, MSC, HSC at the cellular level.

### Application of the model to simulate bone remodeling in mouse vertebrae

The overview of the simulation model can be seen in Figure 1B. Each voxel has a value of bone mineral density from 0 to 1160 mg HA/cm^3^ and this value is converted to grayscale values ranging from 0 to 1 with linear scaling. We considered the grayscale bone density values from 0.5 to 1 as bone tissue. The minimum value of 0.5 corresponds to a bone density value of 580 mg HA/cm^3^ which is the same threshold value employed previously for the postprocessing of the original corresponding *in vivo* data (Scheuren et al. 2020a).

The mechanical signal was obtained using the micro-finite element analysis (micro-FE). For this purpose, we assumed bone tissue to be an isotropic, homogenous material. The bone tissue and marrow voxels of CV6 vertebrae were converted to Young’s modulus values (14.8 GPa and 2 MPa, respectively) and were assigned a Poisson’s ratio of 0.3 as performed previously by Webster et al. (2008). At the proximal and distal ends of the micro-FE model, two cylindrical discs were added to simulate the presence of the intervertebral discs (IVD) (Webster et al. 2008). The nodes at the proximal end of the micro-FE mesh were set to have 0 displacements in all directions, while a displacement of 1% along the longitudinal axis of the micro-FE model was set for the nodes at the distal end. From this analysis, the strain energy density (SED, in MPa) was computed, from which EFF was derived (Pistoia et al. 2002) after rescaling to match the forces applied *in vivo*, which amounts to 4 N for physiological loading in the sham-loaded group (Christen et al. 2012). EFF was used for defining the mechanical signal of the cells (Pistoia et al. 2002; Tourolle et al. 2021) in the single-cell mechanomics: each cell produced an amount of cytokines or unmineralized tissue or resorbed (un)mineralised tissue, following a specific mechanomics curve with a sigmoidal shape to represent the anabolic (osteoid), anti-catabolic (OPG), catabolic (RANKL, osteoid, mineral), anti-anabolic (Scl) response to the mechanical signal (Supplementary Material 1.1). For each cytokine-cell production, the Hill curve was defined for both marrow and bone cells with a specific maximum value, Hill coefficient and *mech*_*thres*_ which is the value of the mechanical signal corresponding to half of the maximum value produced by a cell. These values were defined specimen-specific in order to take into account the variability of the mechanical environment between different animals.

The definition of the Hill parameters was calibrated to have a higher production of cytokines in regions where the local mechanical signal, Gaussian-dilated (sigma 2.5, support 7.5) EFF (ε), is relatively higher or lower for anabolic, anti-catabolic cytokines or catabolic, anti-anabolic cytokines, respectively. The *mech*_*thres*_ for bone cells is defined for every vertebra as follows: 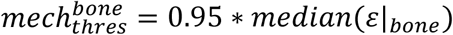 where *ε*|_*bone*_ is the local mechanical signal in the trabecular region of the given vertebra. The mechanomics for the marrow cells that can release products into the volume is defined using this mechanostat threshold: 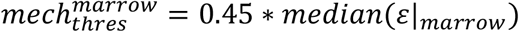 where *ε*|_*marrow*_ is the local mechanical signal in the trabecular region of the marrow of the given vertebra. In particular, *ε*|_*marrow*_ is the gaussian dilated EFF in the trabecular marrow voxels and is used as input of the mechanomics function for the osteoclasts, determining how much bone is resorbed based on the signal. Analogously, *ε*|_*marrow*_ is used as input of the mechanomics function for the osteoblasts and lining cells, determining how much cytokines and osteoid are added to the specific voxel. This sample-based definition was adopted to take into account the intrinsic variability in the strain distribution across different samples due to different trabecular bone microarchitecture. The definitions of 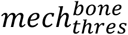 and 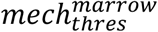 were empirically estimated to discriminate cytokine production in regions of high and low strain relative to the spatial strain distribution of the given sample. Therefore, they were not derived from an estimation of a general mechanostat valid for all samples. We used the mechanical strain to drive the response through regulation of the RANKL-RANK-OPG axis and Scl-LRP5/6 axes which regulate the cell differentiation of the cells. Hence, the formation and resorption responses depend on the strain distribution meaning that in highly loaded regions, formation will be higher than resorption and vice versa.

The model was implemented according to the following multiscale temporal discretization. The cellular behavior described above was simulated with a 40-minute time step (dt_cells-RDD_). Following the simulations obtained by Tourolle (2019), this value turned out to be sufficiently similar to the values observed experimentally about motility and activity of the bone cells. The proteins and other present chemical substances in the simulation were simulated to React, Diffuse, Decay (RDD, Figure 1C) with the same time step for the cells dt_cells-RDD_, subdivided into 10 equal temporal substeps of 4 minutes through Strang splitting (Strang 1968).

Compared to the reference model, a new parallelized approach has been used where the chemical substances and the cells have been subdivided into subdomains. The model computes the spatio-temporal evolution of the cells and signaling pathways and RDD with the timestep dt_cells-RDD_ in parallel across the subdomains. The micro-FE analysis of the vertebra is computed for updating the mechanical signal perceived by the cells with a predefined update interval of 8 hours (dt_micro-FE_), which is higher than dt_cells-RDD_ to simulate a memory mechanism in the perception of the new mechanical signal during the 12 iterations of the cellular behavior until the subsequent micro-FE step.

### Model generation

The trabecular region was automatically obtained for each sample as described previously (C. Marques et al. 2023) comprising a lattice of up to 200×200×300 voxels of the same resolution of *in vivo* data, hence 12 million voxels. In a first step, the greatest connected component (GCC) of the vertebra was defined. The bone phase of the trabecular region (bone mineral density greater than 580 mg HA/cm^3^), was then used for setting the initial mineral and osteoid concentrations to 1. The marrow of the trabecular region was seeded randomly with MSCs and HSCs, with a density of 1.25 × 10^7^ cells/ml for each of these two cell types. Ob and Oc were seeded randomly occupying a portion of the bone surface. The binding sites of the cells were modified to make the multicellular system closer to a real state, where the cells’ receptors are partially or fully bound. Ot were seeded using an exponential distribution to have more osteocytes embedded deeper in bone rather than close to the bone surface. The seeding of all cells in a cross-sectional slice can be seen in Figure 1D. The cytokines were calibrated after running simulations and checking whether the cells would reside and be active in corresponding biochemical regions, e.g. Oc resorb bone and lining cells are present mainly in low strain regions whereas Ob release osteoid primarily in high strain regions. The summary of the initial concentrations is presented in Supplementary Table S2.

The model started running without changes to the bone microarchitecture for 48 iterations (corresponds to 2 days), see Figure 1B, to enable a more adequate spatial arrangement of the cells and cytokines. This pre-processing step is needed because the micro-CT image used as input contains only information regarding the bone microarchitecture. The final configuration after this initialization is illustrated in Figure 1E, where for simplicity reasons only the surface cells, the RANK binding site occupancy for osteoclasts and preosteoclasts and the spatial RANKL distribution in the trabecular region are shown. The model then continued running with changes to bone microarchitecture for five days to enter the active bone remodeling phase and to reduce the dependency of the initial data, where the distributions of the cells, proteins and receptors were affected by uncertainty or absence of such input data. The result of this phase was considered as the initial state for the homeostatic as well as for the simulations where the maximum amount of OPG, RANKL and Scl produced by osteocytes are changed.

### Design of simulations

In this study, we simulated homeostatic remodeling and we performed a one-variable-at-a-time sensitivity analysis of protein expression, where we tested the effects of different production values of RANKL, OPG, and Scl by osteocytes and computed the corresponding static and dynamic bone morphometry data. For each condition different from homeostatic, one parameter was increased or decreased at a single time-point and all the other parameters were kept constant. This approach was used regardless of the simulated animal. Each condition was simulated using *in vivo* data of 5 CV6 vertebrae. The output of the homeostatic condition was compared against the *in vivo* data and against the simulated conditions of higher and lower production values of OPG, RANKL and Scl separately.

First, homeostatic remodeling was simulated with a set of 41 parameters that were kept constant regardless of the simulated animal (Supplementary Table S1). The parameters were optimized for a balanced spatiotemporal evolution of the cytokines, cell differentiation, bone formation and bone resorption. They range from the frequency of random movement of the cells, thresholds regulating the differentiation of the cells, binding site numbers, osteoblast and osteoclast polarization factors, osteoblast and osteoclast cluster size, proliferation and apoptosis rates of cells (especially osteoblasts, MSC, HSC), mechanostat coefficients, the maximum single-cell production rate of cytokines, diffusion and decay coefficients for the cytokines, and mineralization rate. The motility parameters are presented as probability values of movement of 1 voxel per dt_cells-RDD_ in a range of 0 to 1. The Ob and Oc polarization coefficients are values that represent the tendency of these cells to add osteoid or resorb bone, respectively, towards the gradient of the mineral concentration from their position (Tourolle 2019). The competitive reaction RANKL-RANK-OPG requires the definition of the forward and backward binding coefficients for the RANKL-RANK and RANKL-OPG complexes, whereas the simple receptor-ligand kinetics LRP5/6-Scl requires the definition of the forward and backward binding coefficients for the complex LRP5/6-Scl.

To achieve anabolic, anti-anabolic, catabolic and anti-catabolic responses, the variations of the levels of OPG, RANKL and Scl were tested separately by changing the maximum production of OPG, RANKL and Scl by a single Ot 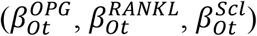 in the simulations using the values reported in Supplementary Table S3, Supplementary Table S4 and Supplementary Table S5. Starting from the baseline value reported in Supplementary Table S1, these higher and lower values were obtained by adding or subtracting the rescaled standard deviation of the serum concentrations levels reported by Shahnazari and colleagues (Shahnazari et al. 2012).

### Morphometry and visualization

Static morphometric parameters analyzed were bone volume fraction (BV/TV), specific bone surface (BS/BV), bone surface density (BS/TV), trabecular thickness (Tb.Th), trabecular spacing (Tb.Sp), and trabecular number (Tb.N). We computed them for each day of the simulation. Dynamic parameters were mineral apposition rate (MAR), mineral resorption rate (MRR), bone formation rate (BFR), bone resorption rate (BRR), mineralizing surface (MS), and eroded surface (ES). They were computed by overlaying the images with a time interval of 2 weeks as analyzed in recent publications (Scheuren et al. 2020a, b), using the image processing language (XIPL, (Hildebrand et al. 1999)). Static parameters between different datasets were compared as percent changes by normalizing the values to the initial value of the morphological parameter of interest while dynamic parameters were compared as absolute values.

The trabecular 3D bone microarchitecture was visualized using ParaView (Kitware, Version 5.10; Clifton Park, NY). The formed, quiescent and resorbed (FQR) regions in the *in vivo* images were obtained after registration of the acquired images and their overlap. The FQR regions in the *in silico* images were directly obtained by overlapping the images at different time points.

### Cluster analysis

Cluster analysis employed the data from the simulation of homeostatic bone remodeling after the selection of a representative animal from the five mice. We determined the formation, quiescence and resorption events in the whole trabecular region by overlapping the last time point of the simulation (t=28 days) against the first time point (t=0 days). A label was assigned to each individual osteocyte based on the closest remodeling event, namely “Formation”, “Resorption” or “Quiescence”. The label “Multiple” was assigned when multiple events had the same distance to the same osteocyte.

In this analysis, effective strain calculated at the first time point was used to classify the levels of mechanical signal where the osteocytes reside. The voxels with mechanical signal higher than the 75^th^ percentile were labeled as “high”, the voxels with mechanical signal lower than the 25^th^ percentile were labeled as “low” and the voxels in between were labeled as “medium”. Then, a representative cross-sectional section with thickness of 5 voxels (52.5 μm) was extracted from the trabecular bone region. The section was selected to show a sufficient amount of bone and spatial variability in effective strain. From this region, the osteocytes, the protein levels, and the mechanical signal in the osteocytes’ positions were used for the subsequent steps.

The cluster analysis adopted in this study employed the protein levels of RANKL, OPG and Scl at the first time point of the simulation. The protein levels were obtained from the 3D positions where the osteocytes reside, similarly to the mechanical signal. The k-means clustering was set with a fixed seed to allow reproducibility; it was applied to the data (Sculley 2010), with a number of clusters ranging from 1 to 10, and a curve with 10 inertia values was obtained. The knee of such curve was used for assigning the clusters to the osteocytes available in the section (Satopää et al. 2011). Cluster labels were assigned to corresponding osteocytes such that each osteocyte has one label for cluster, one for remodeling, and one for mechanical signal.

Using the labels previously assigned for each osteocyte to describe the closest remodeling event, the probability of the remodeling events was computed including the osteocyte distributions within each cluster. Similarly, the probability of high, medium, and low levels of the mechanical signal was computed.

### Software platform

A hybrid C++/Python code (Python Language Reference, Version 3.8) that expanded from the original implementation of the model was used to perform the simulations (Tourolle 2019). The implementation is made available through Python bindings with pybind11 (Jakob et al. 2017). Taking advantage of the other packages used for image processing, analysis and parallelization, we used Python as the front-end. In particular, we used the mpi4py package for managing the MPI parallel distributed computing interface in Python (Dalcin et al. 2011). We employed distributed and shared parallel computing paradigms (OpenMP and MPI) due to the high number of variables used in the simulations. The subdivision in subdomains was carried out to minimize the volume of data from one subdomain to the other (Supplementary material 1.2). We used the MPI parallel version of an algebraic multigrid solver, AMGCL, to solve the diffusion problem of the cytokines into the volume (Demidov 2019). The Swiss National Supercomputing Center (CSCS, Lugano, Switzerland) computational platform (Piz Daint, a Cray XC40/50 model) was used for running the simulations. Each node has 36 cores which can be scheduled to work in a customized way regarding memory and parallelization of tasks. The parallel solver for the spatio-temporal step of the cells and cytokines required 8 nodes to have enough memory and sufficient speedup. For these resources, we use 4 MPI tasks for each node and 9 OpenMP threads. Additionally, the micro-FE analysis of the vertebra models were solved using 2 nodes using ParOsol, a parallel solver designed for micro-FE analysis based on micro-CT images (Flaig and Arbenz 2011). The number of nodes was chosen to have a good trade-off between computational time and speed-up of the code. These two solvers were combined to obtain a suitable environment for solving the interconnected mechanical environment and the tissue, cellular and signaling pathways with a resolution of 10.5 um.

### Statistical analysis

Statistical analysis was performed in R (R Core Team (2019), R Foundation for Statistical Computing, Vienna, Austria). The R lmerTEST package (Kuznetsova et al. 2017) was used to perform the linear mixed model. The linear mixed-effects models account for “intra-correlations” between the simulations and *in vivo* repeated measurements. The model is described in two parts: fixed effects and random effects. The random effects part accounts for the intra-correlations of repeated measures in *in vivo* samples and *in silico* simulations or the high/medium/low production level of OPG, RANKL and Scl. The fixed-effects part accounts for the impact of various covariates over time on outcomes on an average level of the dynamic and static bone morphometric parameters. Furthermore, the likelihood ratio test was performed to assess the goodness of fit of three nested models based on the ratio of their likelihoods. All the nested model equations are in the supplementary section (Supplementary Material 1.3). The significance level between the groups (*in vivo* data/*in silico* simulation or high/low, high/basal, basal/low) was calculated using pairwise comparisons with Tukey’s post-hoc correction for multiple comparisons. The significance level for the interaction between time and group was calculated using repeated measures ANOVA with the linear-mixed model. The mean and the standard error of the mean were plotted and p-values smaller than 0.05 were considered significant (Lenth 2022).

## Results

To study trabecular bone remodeling, we ran our micro-MPA model on five CV6 vertebrae from mice obtained using micro-CT imaging. Using high-performance computing, the computational capability and the efficiency of the computational code were increased, allowing the simulation of thousands of cells and RDD of proteins in a complete trabecular volume. A simulation of 4 weeks took usually 3 to 6 hours on a supercomputer. Longer durations resulted from bigger trabecular volumes (TV) or when the finite-element analysis required more iterations for converging to a numerical solution. A single cell and RDD step took usually up to 10 s, whereas solving a single micro-FE analysis took less than a minute. Our *in silico* micro-MPA model simulated the presence of bone cells and their interaction with the local mechanical environment, the interaction between them and how osteoblasts and osteoclasts add and resorb bone, employing the same isotropic voxel resolution of the 3D micro-CT image. First, we report the results of homeostatic remodeling compared to the *in vivo* data. Then, to demonstrate that the osteocytes can regulate the other cells’ activity, we report the results obtained after the individual manipulation of the maximum production levels of OPG, RANKL and Scl by the osteocytes. The significance achieved for each statistical test is reported in the Supplementary Tables S6-S9.

### Homeostatic remodeling

The simulations were compared to the *in vivo* data of sham loading animals to investigate to which extent single-cell mechanomics of the mechanical signal can simulate homeostasis. Figure 2 shows the *in vivo* and the *in silico* data of homeostatic bone remodeling. In Figure 2A, the individual normalized trabecular bone volume fraction (Norm. BV/TV) is shown for the complete original dataset, illustrating high variability within the group and for each animal over time. Conversely, the *in silico* data of Norm. BV/TV were more stable after the initial remodeling phase. The values of *ε* are shown in Figure 2B with regions of high and low mechanical signal perceived by the osteocytes. *In vivo* data shows that bone formation and bone resorption occur throughout the bone microarchitecture, whereas in the *in silico* results only bone formation occurs more widely in the trabecular region, see Figure 2C. Indeed, bone resorption is more localized at the top of the trabecular region where there are more regions of medium to low mechanical signal compared to the bottom of the trabecular compartment (Figure 2B). In this regard, to emulate the loading condition induced *in vivo*, IVD were added to the *in silico* model and the simulated compressive force was applied to the proximal IVD while fixing the distal IVD. This setting enabled computing a realistic estimation of the mechanical signal (Webster et al. 2008), based on EFF, along the entire trabecular microarchitecture that could be leveraged in the micro-MPA to drive remodeling responses accordingly. Moreover, from visual inspection, bone formation and bone resorption can alternate with each other in the same local region over time *in vivo*, whereas they showed little spatial turnover *in silico*. The static trabecular bone parameters are shown in Figure 2D. BV/TV was not significantly different *in silico* compared to the *in vivo* data on average while it showed significant changes when the interaction with time is considered (n.s. between groups and p<0.01 for the interaction time-group). Tb.Th changed over time in a similar way between the simulated data and the *in vivo* data and these two groups had a similar average (n.s. between groups and for the interaction time-group). While Tb.Sp and Tb.N remained relatively constant over time *in vivo, in silico* we observed a significant decrease and an increase in Tb.Sp and Tb.N, respectively (p<0.05 and p<0.001 between groups, respectively, and p<0.0001 for the interaction time-group). Additionally, we also observed an increase in BS/BV and BS/TV *in silico* but *in vivo* these values remained closer to their initial value (p<0.01 and p<0.05 between groups, respectively, and p<0.0001 for the interaction time-group). MAR obtained from the simulations was significantly lower than the *in vivo* values (p<0.01 between groups) and MRR was much higher *in silico* compared to the *in vivo* data (p<0.001 between groups); the values of MAR and MRR developed differently between the *in silico* and the *in vivo* groups (p<0.001 and p<0.05 for the interaction time-group, respectively). The simulation results captured the BFR average value over time (n.s. between groups) but its temporal evolution was significantly different compared to the *in vivo* data (p<0.01 for the interaction time-group). BRR had similar values *in vivo* and *in silico* (n.s. between groups) with a similar trend over time (n.s. for the interaction time-group). Conversely, MS and ES were lower in the simulations compared to the *in vivo* data (p<0.0001 between groups). MS presented a different temporal progression (p<0.05 for the interaction time-group) while ES did not show a significantly different temporal evolution (n.s. for the interaction time-group).

**Figure 2.**
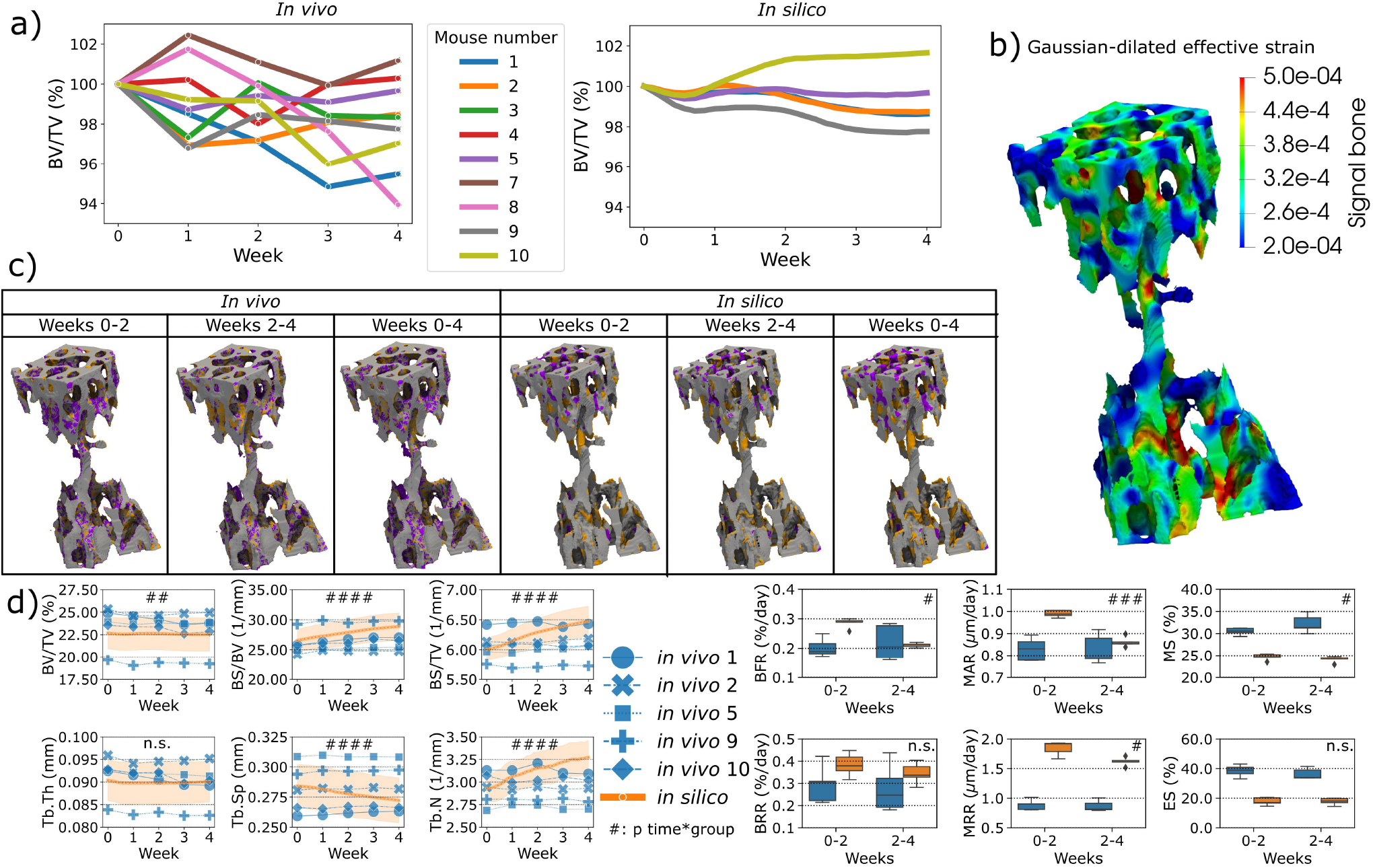
The *in vivo* data and the comparison against the *in silico* modeling of homeostasis. (**A**) The normalized bone volume fraction (Norm. BV/TV) for the complete group of mice *in vivo* (left, n=10) and for the group of mice analyzed *in silico* (right, n=5). (**B**) Mechanical signal in the trabecular region for the representative mouse (number 5). (**C**) Bone formation, quiescent and resorption events images over time *in vivo* and *in silico*. (**D**) Static and dynamic bone morphometry values for the *in vivo* and *in silico* groups. In these plots, only the same selection of mice was plotted for both the *in vivo* and *in silico* data (n=5). (# p<0.05, ## p<0.01, ### p<0.001, #### p<0.0001 for the interaction time*group determined by two-way ANOVA).

### Single-cell mechanomics cluster analysis

In total, 3571 Ot were analyzed from one trabecular cross-section of a mouse caudal vertebra, where Figure 3A-C shows Ot and their associated protein levels of RANKL, Scl and OPG embedded within the local mechanical environment of trabecular bone, respectively. From these sections, higher values of RANKL and Scl were observed in regions with lower EFF whereas higher values of OPG were found in regions of higher EFF. The frequency of the high, medium, and low mechanical signal associated with Ot is shown in Figure 3D. The total probability of labeling high, medium, or low mechanical signal in Cluster 1 was higher compared to Cluster 2 and 3 due to the high number of Ot in Cluster 1. In Cluster 1, the highest probability of high mechanical signal within the cluster was detected, while the highest probability of low mechanical signal within the cluster was detected in Cluster 2. The frequency of the closest bone remodeling events to Ot is shown in Figure 3E. Similar to Figure 3D, the probability of closest bone remodeling in Cluster 1 was higher compared to Clusters 2 and 3 due to the high number of Ot in Cluster 1. The highest probability of bone formation, bone resorption and bone quiescence was observed in Cluster 1. Intermediate and lowest probabilities of bone formation and resorption in Cluster 3 were obtained, respectively. Conversely, intermediate and lowest probabilities of bone resorption and formation were obtained in Cluster 2, respectively.

**Figure 3.**
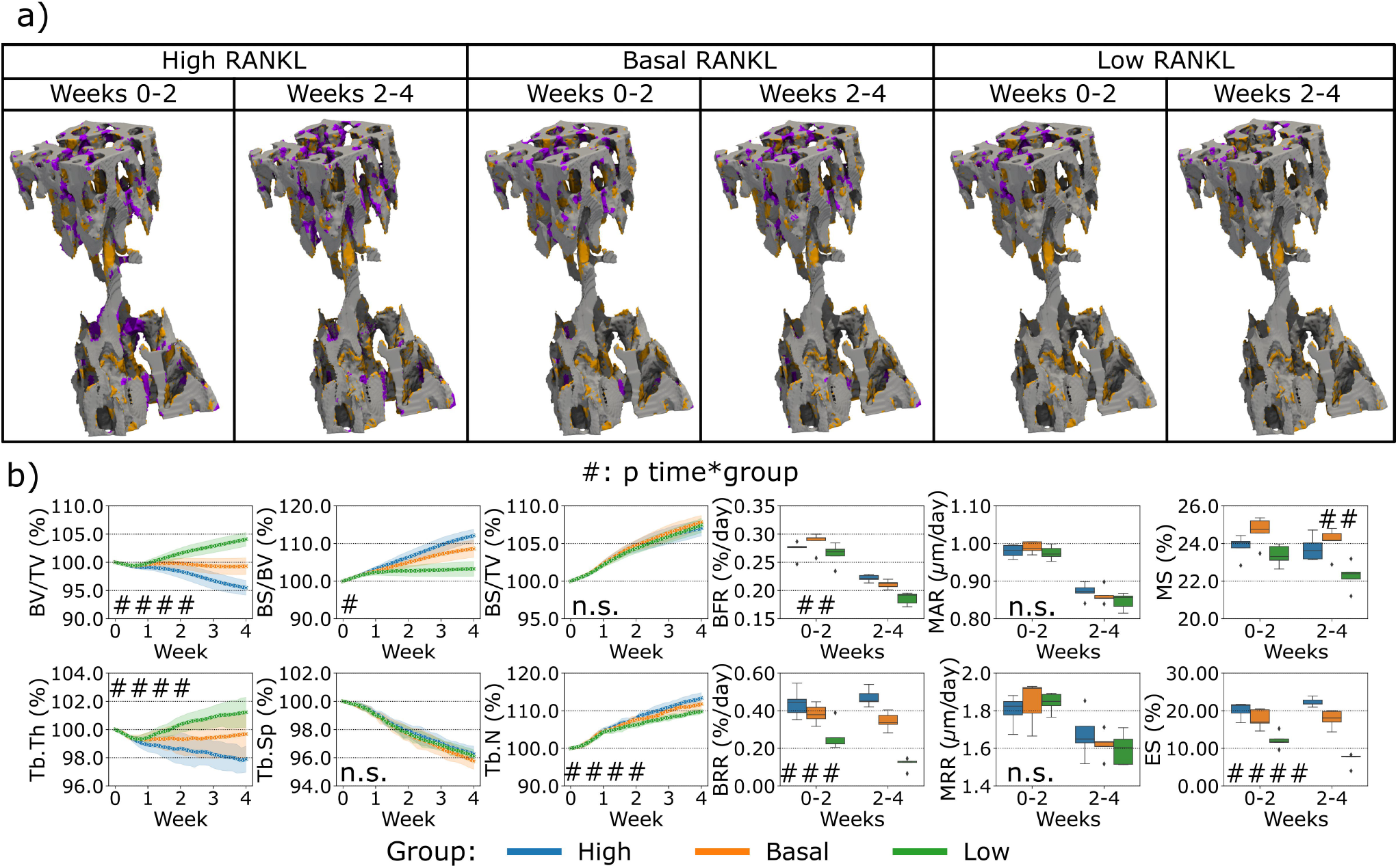
Cross-section of trabecular bone with osteocytes and their protein levels, mechanical environment and bone remodeling from the simulation of homeostatic bone remodeling using single-cell mechanomics. (A), (B), and (C) show trabecular bone colored with the mechanical signal (EFF=effective strain) and osteocytes (as dots, n=748) with Receptor activator of nuclear factor *κ*β ligand, sclerostin (Scl) and osteoprotegerin (OPG) levels, respectively. RANKL, Scl and OPG are shown in shades of reds, greens and blues, respectively. (D) Absolute frequency of levels of mechanical signal in the section divided per cluster. Yellow=high label, gray= medium label, light blue= low label. (E) Absolute frequency of closest bone remodeling events (Formation, Quiescence, Resorption or Multiple if more than an event is closest) to the osteocytes in the section divided per cluster.

Using the *in silico* tools, we obtained a trabecular bone section where bone remodeling events can be shown along with Ot present in the first time point, see Figure 3F, color-labeled with the cluster number. Some Ot were present in resorbed bone, therefore in the last time point of the simulation they were no longer present. On the other hand, bone resorption was observed to remove some connections in the trabecular microarchitecture whereas bone formation tended to deposit bone in layers. The clustering procedure classified Ot based on the protein levels but not relative to their positions. Therefore, Ot of the same cluster may be found in two or more different regions in the section. For example, Ot in Cluster 3 were found in 8 different locations. Nonetheless, the majority of Ot were classified as a “connected” group.

### Sensitivity analysis of protein expression

First, the variation of OPG levels produced by the osteocytes led to a change in bone remodeling activity from the cells, see Figure 4. The main effect of the spatial characterization of the remodeling regions was a higher or lower catabolic activity in the distal end of the CV6 when OPG was reduced or increased respectively, see Figure 4A. This cell activity was reflected in the Norm. BV/TV with lower values when OPG was lower and higher values when OPG was higher (p<0.01 for the group comparison high OPG-basal OPG, basal OPG-low OPG and p<0.0001 for the interaction time-group), see Figure 4B. Norm. Tb.Th. showed no significant difference for the interaction time-group, however the values computed with lower OPG were significantly lower than the values using high and basal OPG levels (p<0.05 for the group comparison high OPG-low OPG and basal OPG-low OPG). Norm. BS/BV showed an inverse relationship with the OPG levels (p < 0.01 and p<0.05 for the group comparison high OPG-basal OPG and basal OPG-low OPG, respectively). In addition, Norm. BS/BV showed different curves over time (p<0.05 for the interaction time-group). Norm. BS/TV, Norm. Tb.Sp and Norm. Tb.N were not affected by the changes in OPG levels (n.s. between groups and for the interaction time-group). BFR was not significantly affected by the variations in OPG with a similar slightly decreasing trend over time among the three levels. The BRR and ES showed a similar variability of Norm. BS/BV, with lower values when OPG was higher and higher values when OPG was lower (p<0.01 for the group comparison high OPG-low OPG and p<0.05 for the other group comparisons). The significance of these changes was observed for both BRR and ES over time between groups data (p<0.05 for the interaction time-group). MS showed significantly different values between groups (p<0.05), however only the comparisons of high OPG-basal OPG and basal OPG-low OPG showed significant differences between the groups (p<0.01 and p<0.05, respectively). MAR and MRR did not show differences among the groups when the OPG production level was changed in the simulations. These results suggest that OPG produced by the osteocytes can inhibit the osteoclasts recruitment and the consequent amount of resorption by the available osteoclasts. In addition, the number of osteoclasts also influenced the number of active osteoblasts, as shown by the variability of the MS. The net effect of these changes has an impact on the static parameters, primarily on Norm. BV/TV where the homeostatic balance is lost with alterations of the OPG levels.

**Figure 4.**
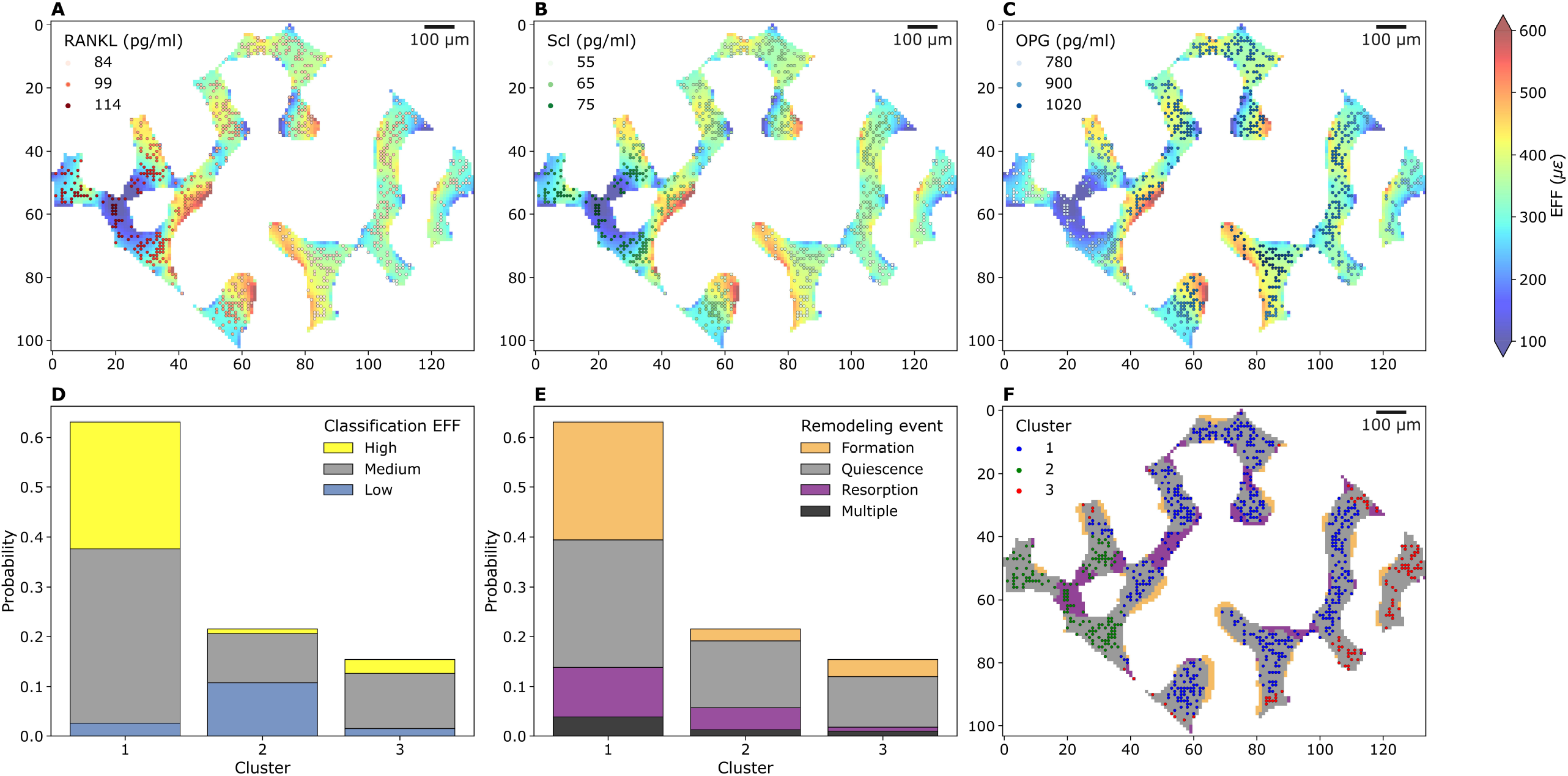
*In silico* results of variations of the maximum single cell production level of osteoprotegerin (OPG) by osteocytes. (**A**) Biweekly FQR regions of results obtained with higher and lower OPG production levels for the representative mouse (number 5. (**B**) Static and dynamic bone morphometry parameters of the results of the simulations of the same group of mice (n=5) under three different production levels of OPG, starting from the same initial condition of the basal level. High=higher production level of OPG, Basal=basal production level of OPG as in the homeostatic configuration, low=lower production level of OPG. (# p<0.05, ## p<0.01, ### p<0.001, #### p<0.0001 for the interaction time*group determined by two-way ANOVA).

Second, the variation of RANKL levels produced by the osteocytes led to a different bone remodeling activity from the osteoblasts and osteoclasts, see Figure 5. Similar to the variations observed for OPG, we observed primarily a lower or higher catabolic activity when RANKL in the distal end of the CV6 was reduced or increased respectively, see Figure 5A. This cell activity was reflected in the Norm. BV/TV, Norm. Tb.Th. with lower values when RANKL was higher and higher values when RANKL was lower (p<0.001 for the group comparison high RANKL-low RANKL, p<0.01 and p<0.05 for the group comparison high RANKL-basal RANKL respectively, p<0.01 and n.s. for the group comparison basal RANKL-low RANKL respectively, p<0.0001 for the interaction time-group), see Figure 5B. Norm. BS/TV did not respond differently to higher or lower RANKL values over time and the average values over time were not statistically significantly different (n.s. between groups and for the interaction time-group). Norm. BS/BV, Norm. Tb.N and BRR and ES were lower when RANKL was lower and they were higher when RANKL was higher (p<0.05 between groups, for all possible group comparisons and for the interaction time-group; detailed significances in Supplementary Table S8). Norm. Tb.Sp was not affected by the changes in RANKL over time and its average value was not affected (n.s. between groups and for the interaction time-group). The changes in RANKL had an impact on BFR, with higher or lower values when RANKL was higher or lower, respectively (p<0.01 between groups and for the interaction time-group). The only comparison which did not show a significant difference in BFR was between high and basal levels of RANKL. BRR was also affected by the changes in RANKL in a similar way (p<0.001 for the group comparison high RANKL-low RANKL and basal RANKL-low RANKL, p<0.05 for the group comparison high RANKL-basal RANKL, and p<0.001 for the interaction time-group). MAR and MRR did not show differences among the groups with changing RANKL production levels. These results suggest that RANKL promoted the osteoclasts differentiation by the changes of ES which in turn led to changes in resorption. The changes in the Ob recruitment are reflected in the extent of the formed surface by the Ob. The change in the formed surface had a cumulative effect on different values of BFR. The net effect of these changes had an impact on the static parameters, with the parameters starting to differ from the basal condition earlier compared to OPG, thus augmenting the separation from the homeostatic condition.

**Figure 5.**
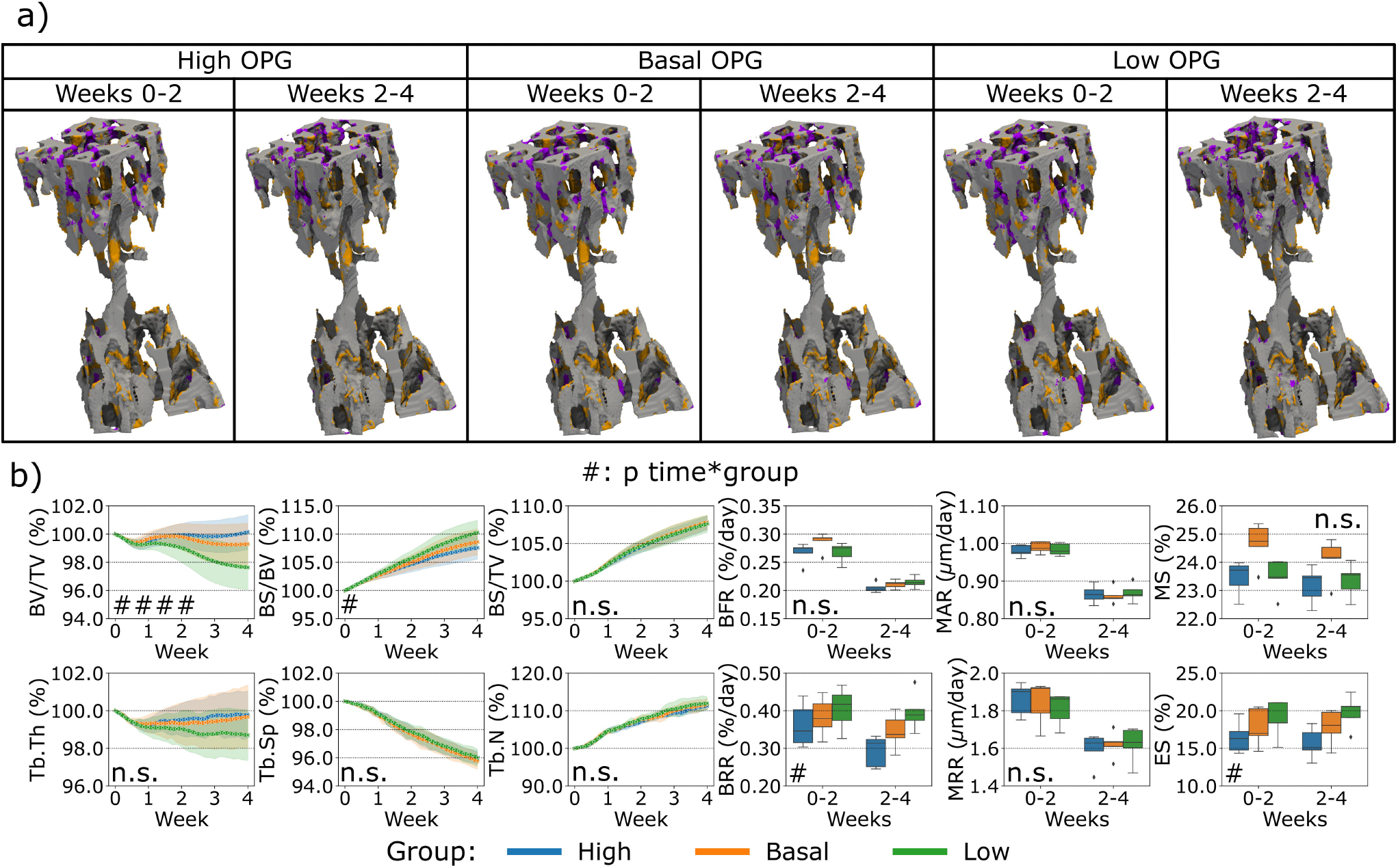
*In silico* results of variations of the maximum single cell production level of receptor activator of nuclear factor kB ligand (RANKL) by osteocytes. (**A**) Biweekly FQR regions of results obtained with higher and lower RANKL production levels for the representative mouse (number 5). (**B**) Static and dynamic bone morphometry parameters of the results of the simulations of the same group of mice (n=5) under three different production levels of RANKL, starting from the same initial condition of the basal level. High=higher production level of RANKL, Basal=basal production level of RANKL as in the homeostatic configuration, low=lower production level of RANK. (# p<0.05, ## p<0.01, ### p<0.001, #### p<0.0001 for the interaction time*group determined by two-way ANOVA).

Finally, the variation of Scl levels produced by the osteocytes led to a different bone remodeling activity from the osteoblasts and osteoclasts compared to what was observed for the variations of RANKL and OPG before, see Figure 6. In the proximal end of the CV6 we observed primarily higher or lower anabolic activity of the osteoblasts when Scl was reduced or increased respectively, see Figure 6A. Therefore, bone formation was affected, and a similar effect was observed in the Norm. BV/TV with lower values when Scl was higher, see Figure 6B. Indeed, the variations of Scl were statistically significant for Norm. BV/TV when comparing high and low Scl values as well as high and basal Scl values (p<0.01). However, their curves were not significantly different over time (n.s. for the interaction time-group). The statistical findings for Norm. BV/TV also apply to Norm. Tb.Th which slightly decreased over time and stabilized afterwards with a high Scl level whereas it increased when Scl was basal or lower (p<0.05 for the group comparisons high-low and high-basal). Conversely, Norm. BS/BV and Norm. BS/TV increased over time. The statistical significance of the variations for Norm BS/BV was the same as of Norm. BV/TV (except for p<0.05 for the group comparison high Scl-basal Scl), whereas Norm BS/TV showed differences only in the group comparison high-basal (p<0.05). Interestingly, no increase in the Norm. BV/TV was observed for lower values of Scl. The temporal evolution of Norm. Tb.Sp and Norm. Tb.N did not present significant differences between the variations of Scl (n.s. for the interaction time-group); Norm. Tb.N was not affected on average by different levels of Scl (n.s. between groups) while Norm. Tb. Sp was only affected by higher values of Scl when compared to basal values of Scl (p<0.05 for the group comparison high Scl-basal Scl). BFR showed a significant variability due to Scl (p<0.05 between groups) but only in the comparisons high-low and high-basal. Also, MS was affected by the variations of Scl in a more evident way (p<0.01 for the group comparison high Scl-basal Scl and p<0.05 for the other group comparisons). Scl did not change how bone was deposited by osteoblasts and this outcome was visible from the absence of variability in MAR with different Scl values (n.s. between groups and for the interaction time-group). BRR, MRR and ES did not show differences when Scl levels changed, nor its effect was significantly visible over time (n.s. between groups and for the interaction time-group). The changes in Scl levels had a net effect on the Norm. BV/TV less visible compared to the changes in OPG and RANKL levels, with a temporal separation from the homeostatic range of values similar to the OPG case.

**Figure 6.**
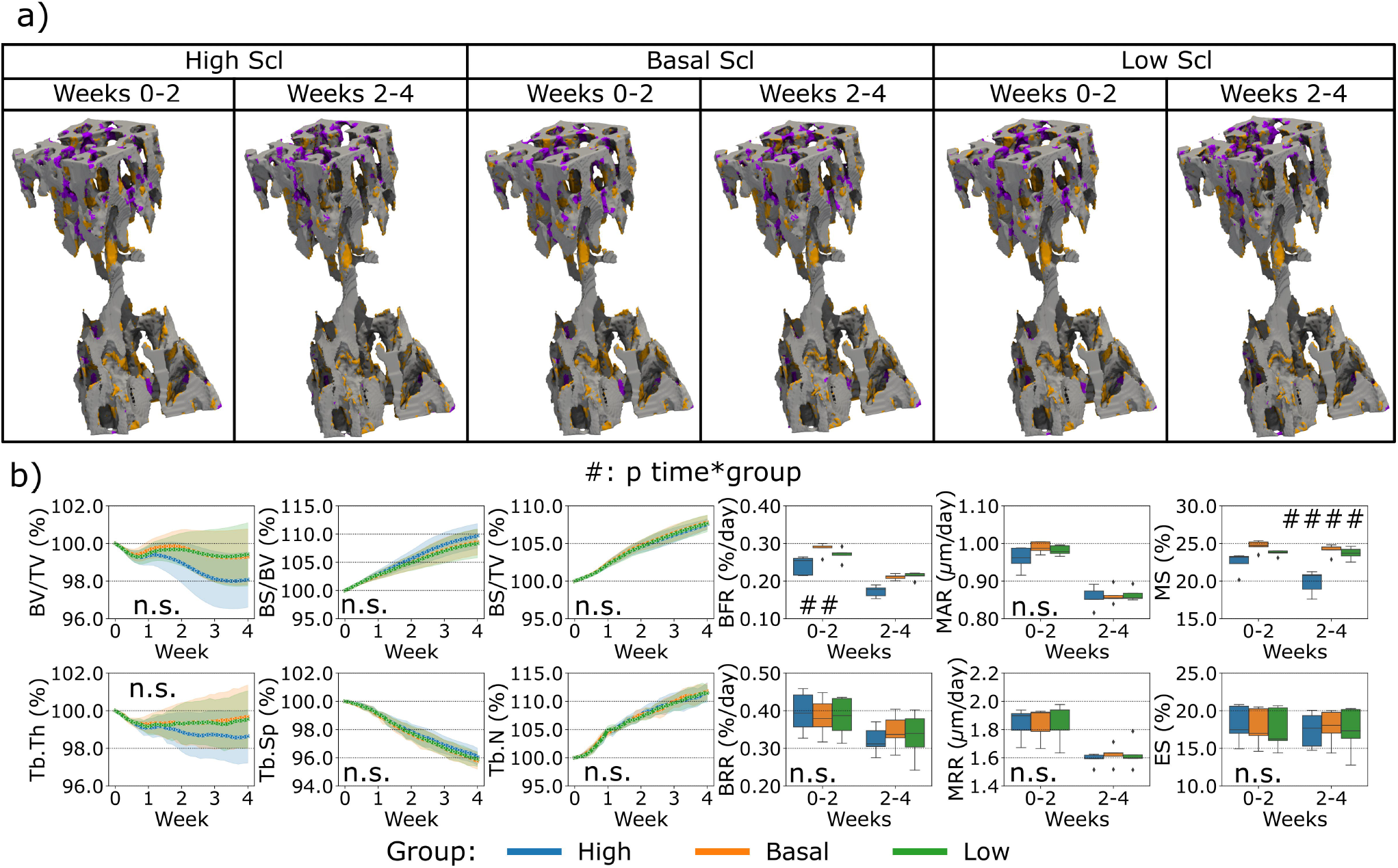
*In silico* results of variations of the maximum single cell production level of sclerostin (Scl) by osteocytes. (**A**) Biweekly FQR regions of results obtained with higher and lower Scl production levels for the representative mouse (number 5). (**B**) Static and dynamic bone morphometry parameters of the results of the simulations of the same group of mice (n=5) under three different production levels of Scl, starting from the same initial condition of the basal level. High=higher production level of Scl, Basal=basal production level of Scl as in the homeostatic configuration, low=lower production level of Scl. (# p<0.05, ## p<0.01, ### p<0.001, #### p<0.0001 for the interaction time*group determined by two-way ANOVA).

## Discussion

The proposed *in silico* micro-MPA model successfully simulated trabecular bone remodeling to evaluate the static and dynamic morphometry of the trabecular bone microarchitecture. We demonstrated that our model can simulate a homeostatic response similar to that observed with *in vivo* data. Although some morphological aspects were not captured by the simulations, our model showed adaptation towards a normalized trabecular bone fraction in all samples. Furthermore, the variability observed *in vivo* was also partially reflected in our simulations. Nonetheless, these differences must be understood in the context of a novel multiscale micro-MPA model of trabecular bone remodeling that demonstrates for the first time physiological behavior of the cells, where group averages of key static morphometric parameters (trabecular bone volume fraction and thickness) follow *in vivo* data. Capturing the static parameters with *in silico* models has been proven to be challenging and the most frequent parameter to be captured was BV/TV (Schulte et al. 2013b; Levchuk et al. 2014; Levchuk 2015). This parameter did not show significant differences on average between the *in silico* and the *in vivo* values, whereas Tb.Sp and Tb.N were different in our simulations compared to the *in vivo* data. Our simulations showed a similar evolution pattern of the static bone morphometric parameters across the samples in the second half of the simulation, meaning that it was capable to simulate similar bone changes for a group of samples with limited variability in the output. The single-cell activity and the signaling pathways were additional information modeled and they could be further tuned to capture the spatiotemporal evolution of the bone microarchitectures. With this model, it is possible to test hypothesis or translate findings from other studies to improve the fidelity and the match against experimental data, e.g. considering fluid flow inside the lacunocanalicular network improves prediction of bone remodeling (van Tol et al. 2020). This study might be considered a step further in modeling the action of single cells and the signaling pathways compared to models of bone remodeling based on systemic ordinary differential equations (ODEs) where the spatial information of the cells is missing (Buenzli et al. 2012; Pastrama et al. 2018; Martin et al. 2019). With this model, it will be possible to quantitatively investigate the mechanobiological properties at the cellular and protein levels that are associated with ageing in mice and in the future to use the 3R principles to replace, reduce and refine such animal models.

Tb.N, Tb.Th and Tb.Sp could be dependent on the single-cell movement, apposition and resorption rates by osteoblasts and osteoclasts as well as by the mineralizing and eroded surface. The single-cell movement was modeled with migration towards regions of higher or lower strains for the osteoblasts and osteoclasts, respectively. Such migration can lead to bone formation and bone resorption patterns that can alter the morphology of the microarchitecture and the extent of the remodeled surface to adapt to the local perceived strains. Further, the mineral apposition and resorption rates had to be higher to reduce the difference between *in silico* and *in vivo* bone formation and resorption rates. Eventually, trabecular parameters would be affected by these changes resulting from more accentuated local remodeling. If MAR, MRR, MS and ES were closer to the *in vivo* values, the static trabecular parameters could have better followed the corresponding *in vivo* values. However, this aspect should be further inspected because, for example, the *in silico* model of Levchuk and colleagues captured Tb.N, Tb.Th and Tb.Sp despite a significant discrepancy in dynamic parameters (Levchuk et al. 2014), while the model of Schulte and colleagues for ovariectomy and loading captured some dynamic parameters and BV/TV but not Tb.N, Tb.Th and Tb.Sp (Schulte et al. 2013b). In the model presented, the regulation of the dynamic parameters was not trivial because the mineralizing and eroded surfaces were not easy to capture due to the complex interplay between the mechanical environment, the mechanomics and the subsequent cascade in the signaling process to regulate osteoblasts and osteoclasts activity. The remodeled regions were rather large *in silico* compared to *in vivo* which were more scattered throughout the trabecular volume. Thus, the trabecular remodeling by the single cells was less localized *in silico*.

Our proposed cluster analysis of the osteocytes was different compared to previous experimental approaches, where the osteocytes were obtained only at the end of the study (Trüssel 2015; Paul 2020; Scheuren 2020). In those approaches, the backward correlation of bone remodeling events with the final osteocyte population can show only the final distribution of the osteocytes. Conversely, the clustering analysis applied here allowed the selection of a section of interest and the study of the osteocyte population available at the beginning of the study. The clustering analysis identified one cluster (Cluster 1) where the cells showed a higher probability of high mechanical signal and bone formation events. Cluster 2 was characterized by higher levels of RANKL and Scl, a higher probability of low mechanical signal and higher probability of bone resorption within the cluster. Taken together, these findings confirmed the regulation of the pathways from tissue to cell scales, in alignment with the current observations that high mechanical signal is associated with lower expression of RANKL and Scl (Robling et al. 2008; Galea et al. 2020), higher expression of OPG (You et al. 2008) and higher bone formation (Schulte et al. 2013a).

We demonstrated OPG inhibits excessive osteoclastic bone resorption as it was observed previously (Kramer et al. 2010; Cawley et al. 2020). Cawley et al. (2020) showed OPG to be secreted by osteoblasts whereas Kramer et al. (2010) found osteocytes to be the cells mainly producing OPG. Our analysis highlighted osteocytes are the cells responsible for the production of RANKL and the subsequent recruitment of osteoclasts and increase of bone resorption, as it was shown in previous studies (Nakashima et al. 2011; Xiong et al. 2015). In our study, osteocytes were also identified as sources of sclerostin inhibiting the osteoblastic activity and therefore reducing bone formation. This finding was experimentally obtained also by Van Bezooijen et al. (2004), where sclerostin protein was found to be expressed by osteocytes and not by osteoclasts in cortical and trabecular bone. Further, they also observed the inhibitory effect of sclerostin on osteoblasts, confirming its importance for regulating bone formation (Winkler et al. 2003; Li et al. 2008; Colucci et al. 2011).

We observed that affecting the catabolic response would also imply a change in the anabolic activity. This aspect was evident from the variations of RANKL and OPG which had a direct impact on the osteoclasts’ activity and subsequently affecting the osteoblasts’ activity. Besides, the anti-anabolic response would also imply a subsequent change in the catabolic activity, as it was seen from the variations of Scl which inhibited osteoblasts and a consequent slower resorption activity. The variations in the RANKL, OPG and Scl did not lead to proportional changes in MS because of the dynamic equilibrium the cells can reach, meaning that the levels of cytokines can result in a non-linear response in the amount of mineralized surface. These findings agree with the coupling between bone formation and bone resorption events as observed *in vivo*. Our *in silico* model represented bone formation mostly occurring over time in the same region where the strains were high, until the osteoblasts de-differentiated into lining cells or became embedded into the osteoid and differentiate into preosteocytes. On the other hand, bone resorption started from regions where strains were low, which also had higher RANKL, leading to more prominent recruiting of osteoclasts. TGF-β1 is thought to be responsible for coupling bone resorption and bone formation (Raggatt and Partridge 2010; Kasagi and Chen 2013; Weivoda et al. 2016; Durdan et al. 2022) and osteoclasts resorb bone in regions of low strains. MSCs proliferated closer to osteoclasts due to the presence of TGF-β1 being released upon resorption in these regions of low strains. In addition, in such regions, sclerostin was also produced by osteocytes to inhibit bone formation by the osteoblasts. Therefore, the osteoblastic differentiation of such cells was inhibited and MSCs would directly differentiate into lining cells instead of osteoblasts in these regions of low strains.

It is not yet fully understood when the osteoclasts stop resorbing bone (Filgueira 2010), but it is assumed that this happens when the mineral phase of the bone underneath is degraded to a certain extent (Kanehisa and Heersche 1988). Currently, this mechanism is not reproduced directly in the *in silico* model. However, our model can produce realistic osteoclast cell behavior. These cells have been shown to alternate bone resorption and migration on the bone surface known as pit mode (Delaisse et al. 2021). In our *in silico* model, osteoclasts stopped being active, and therefore resorptive, when they were not in a cluster or if they died by apoptosis or due to lack of bound RANK receptors. Further, the resorption rate of mineral and collagen by osteoclasts was considered the same. This is the case when osteoclasts resorb bone while moving on the bone surface known as trench mode (Delaisse et al. 2021). Conversely, these rates are usually different when osteoclasts resorb bone in pit mode, with solubilization of mineral being faster than the evacuation of collagen fragments (Delaisse et al. 2021). The modelling of these events might be an oversimplification of how osteoclasts stop being active and it does not represent the same frequency of interruption of bone resorption by osteoclasts.

On a different note, our model did not investigate cortical remodeling because it was shown that its mechanical control is different from the control in the trabecular region (Pearson and Lieberman 2004) and their mineralization and turnover rates are different (Lerebours et al. 2020), hence the model could not capture the changes in cortical bone using the same rules and parameters for trabecular bone.

For initializing the cytokines in the model, we used values measured from experimental studies which are closely in line with the modelled environment. However, it is likely that the values appropriate for the mouse vertebra at a given age might differ from the experimental values due to site, age, loading and phenotypic characterization of the data. This problem has been identified for *in silico* models of bone mechanobiology (Checa and Prendergast 2009), where simple approaches were adopted to overcome this lack of information (Perier-Metz et al. 2020, 2022; Borgiani et al. 2021). This limitation might be partially overcome by taking advantage of the mechanical environment as it was performed by Tourolle et al. (2021). A more synergistic study where the cytokines are experimentally obtained from the same site on a regular basis would help in the calibration of this micro-MPA *in silico* model. In addition, the information on the cell densities is scarce and they might change between different bones, age, sex, and physiological conditions. Moreover, the cells might be distributed differently in bone and in the marrow. This aspect is especially relevant for osteocytes which are not motile in bone, meaning that their initial distribution is essentially preserved throughout the time of the simulation, except when they are no longer present due to cell apoptosis or bone resorption. The initial cell densities and cytokine distributions will have implications in the estimations of the proliferation and cell rates as well as the single-cell activity, e.g., resorption and cytokine production rates. Consequently, the amount of unknown initial conditions is very high, and it can take some time before the simulated cells and cytokines can reach a realistic spatio-temporal distribution, with minimal influence of the assumed initial condition. This limitation was addressed by the additional iterations of the model first without changes and then with changes to the bone microarchitecture. However, the number of iterations required for this purpose could be even higher.

The receptor-ligand kinetics in the context of micro-MPA *in silico* models is still very novel and needs further exploration. Depending on the application, the formulations might differ to include other aspects like trafficking and intracellular signaling (Cilfone et al. 2015). Moreover, there might be molecular aspects that might be lost when using this kind of equation and coefficients might be recalibrated depending on the environment. Indeed, some coefficients might be experimentally obtained from analysis of the interactions between monomers (Nelson et al. 2012), but their usage might not be straightforward due to the coexistence of other phenomena like spatial diffusion and movement of the ligands from one receptor to the other in the proximity of the cell surface (Erbaş et al. 2019). Furthermore, the cell response is usually achieved when most of its surface receptors are still unoccupied (Lodish et al. 1999). It is still difficult to estimate *in vivo* the number of surface receptors and the number of occupied receptors along with an accurate description of the receptor-ligand kinetics, especially when competitive reactions are involved, e.g., the RANKL-RANK-OPG axis. Hence, the receptor-ligand kinetics in bone remodeling remains challenging but crucial. Nonetheless, it is our belief that the sensitivity analysis of protein expression presented can encourage more complex approaches such as uncertainty quantification, for calibration and estimation of more realistic model parameters (Geris and Gomez-Cabrero 2016; Gomez-Cabrero et al. 2016; Viceconti et al. 2019; Hamis et al. 2021). Ultimately, we expect these could have direct benefits in the quality of the simulation output.

## Conclusion

This study showcases the use of single-cell mechanomics in a micro-MPA *in silico* model of trabecular bone remodeling applied to *in vivo* data. We were able to reproduce homeostatic bone remodeling and highlight how tuning the single-cell osteocyte production rates of OPG, RANKL and Scl further induces anabolic, anti-anabolic, catabolic and anti-catabolic responses from the baseline model. Specifically, OPG and RANKL levels altered the homeostatic remodeling around the equilibrium value of the average trabecular bone volume fraction while Scl altered the final trabecular bone volume fraction at equilibrium. By careful calibration of biological parameters, we hope that this model can shed light on bone remodeling and associated diseased states. Micro-MPA models over several scales will be needed in the future because only with them it will be possible to unravel biological processes and their effects realistically. This will advance the field of bone remodeling and our current understanding of its mechanisms.

## Author Contributions

DB, FM, CL, JK, YDB, FS and RM contributed to the design of the study. DAB, FM, CL and JK contributed to the development of the model, assembly and assessment of data, AS and FS contributed to the statistical analysis and interpretation of results, DB wrote the main manuscript text. All authors contributed to revising the manuscript and approved the final version to be submitted.

## Acknowledgments

This study was supported by a grant from the Swiss National Supercomputing Centre (CSCS) under project ID s1070.

## Supplementary Material

See additional file for Supplementary Material.

## Data Availability Statement

The raw data supporting the conclusions of this article will be made available by the authors, without undue reservation.

## References

Bezooijen RL Van, Roelen BA, Visser A, et al. 2004. Sclerostin is an osteocyte-expressed negative regulator of bone formation, but not a classical BMP antagonist. The Journal of experimental medicine 199: 805–14.

Boaretti D, Betts DC, and Müller R. 2018. Studying how the link between mechanical stimulation and cellular activation effects bone microarchitecture. In: Book of Abstracts of the 25th Congress of the European Society of Biomechanics (ESB 2019).

Boaretti D, Wehrle E, Bansod Y, et al. 2022. Perspectives on in silico bone mechanobiology: computational modelling of multicellular systems. European Cells and Materials 44: 56–73.

Borgiani E, Duda GN, Willie BM, and Checa S. 2021. Bone morphogenetic protein 2-induced cellular chemotaxis drives tissue patterning during critical-sized bone defect healing: an in silico study. Biomechanics and Modeling in Mechanobiology: 1–18.

Bourhis E, Wang W, Tam C, et al. 2011. Wnt antagonists bind through a short peptide to the first β-propeller domain of LRP5/6. Structure 19: 1433–42.

Boyce BF and Xing L. 2008. Functions of RANKL/RANK/OPG in bone modeling and remodeling. Archives of biochemistry and biophysics 473: 139–46.

Buenzli PR, Pivonka P, Gardiner BS, and Smith DW. 2012. Modelling the anabolic response of bone using a cell population model. Journal of theoretical biology 307: 42–52.

C. Marques F, Boaretti D, Walle M, et al. 2023. Mechanostat parameters estimated from time-lapsed in vivo micro-computed tomography data of mechanically driven bone adaptation are logarithmically dependent on loading frequency. bioRxiv.

Cawley KM, Bustamante-Gomez NC, Guha AG, et al. 2020. Local production of osteoprotegerin by osteoblasts suppresses bone resorption. Cell Rep 32: 108052.

Checa S and Prendergast PJ. 2009. A mechanobiological model for tissue differentiation that includes angiogenesis: a lattice-based modeling approach. Annals of biomedical engineering 37: 129–45.

Christen P, Rietbergen B van, Lambers FM, et al. 2012. Bone morphology allows estimation of loading history in a murine model of bone adaptation. Biomechanics and Modeling in Mechanobiology 11: 483–92.

Cilfone NA, Kirschner DE, and Linderman JJ. 2015. Strategies for efficient numerical implementation of hybrid multi-scale agent-based models to describe biological systems. Cellular and molecular bioengineering 8: 119–36.

Colucci S, Brunetti G, Oranger A, et al. 2011. Myeloma cells suppress osteoblasts through sclerostin secretion. Blood Cancer J 1: e27–e27.

Dalcin LD, Paz RR, Kler PA, and Cosimo A. 2011. Parallel distributed computing using Python. Adv Water Resour 34: 1124–39.

Delaisse J-M, Søe K, Andersen TL, et al. 2021. The mechanism switching the osteoclast from short to long duration bone resorption. Frontiers in Cell and Developmental Biology 9: 555.

Demidov D. 2019. AMGCL: An efficient, flexible, and extensible algebraic multigrid implementation. Lobachevskii Journal of Mathematics 40: 535–46.

Dobson PF, Dennis EP, Hipps D, et al. 2020. Mitochondrial dysfunction impairs osteogenesis, increases osteoclast activity, and accelerates age related bone loss. Sci Rep 10: 1–14.

Durdan MM, Azaria RD, and Weivoda MM. 2022. Novel insights into the coupling of osteoclasts and resorption to bone formation. Semin Cell Dev Biol.

Elson A, Anuj A, Barnea-Zohar M, and Reuven N. 2022. The origins and formation of boneresorbing osteoclasts. Bone: 116538.

Erbaş A, La Cruz MO De, and Marko JF. 2019. Receptor-ligand rebinding kinetics in confinement. Biophysical journal 116: 1609–24.

Filgueira L. 2010. Osteoclast differentiation and function. In: Bone Cancer. Elsevier.

Flaig C and Arbenz P. 2011. A scalable memory efficient multigrid solver for micro-finite element analyses based on CT images. Parallel Computing 37: 846–54.

Franz-Odendaal TA, Hall BK, and Witten PE. 2006. Buried alive: how osteoblasts become osteocytes. Developmental dynamics: an official publication of the American Association of Anatomists 235: 176–90.

Galea GL, Paradise CR, Meakin LB, et al. 2020. Mechanical strain-mediated reduction in RANKL expression is associated with RUNX2 and BRD2. Gene 763: 100027.

Geris L and Gomez-Cabrero D. 2016. An introduction to uncertainty in the development of computational models of biological processes. In: Uncertainty in Biology. Springer.

Gomez-Cabrero D, Tegnér J, and Geris L. 2016. Computational modeling under uncertainty: challenges and opportunities. In: Uncertainty in Biology: A Computational Modeling Approach. Springer.

Hamis S, Stratiev S, and Powathil GG. 2021. Uncertainty and sensitivity analyses methods for agentbased mathematical models: An introductory review. THE PHYSICS OF CANCER: Research Advances: 1–37.

Hildebrand T, Laib A, Müller R, et al. 1999. Direct three-dimensional morphometric analysis of human cancellous bone: microstructural data from spine, femur, iliac crest, and calcaneus. Journal of bone and mineral research 14: 1167–74.

Jakob W, Rhinelander J, and Moldovan D. 2017. pybind11 - Seamless operability between C++11 and Python.

Kanehisa J and Heersche J. 1988. Osteoclastic bone resorption: in vitro analysis of the rate of resorption and migration of individual osteoclasts. Bone 9: 73–9.

Kanis JA, Norton N, Harvey NC, et al. 2021. SCOPE 2021: a new scorecard for osteoporosis in Europe. Archives of osteoporosis 16: 1–82.

Kasagi S and Chen W. 2013. TGF-beta1 on osteoimmunology and the bone component cells. Cell & Bioscience 3: 1–7.

Klein-Nulend J, Bacabac R, and Bakker A. 2012. Mechanical loading and how it affects bone cells: the role of the osteocyte cytoskeleton in maintaining our skeleton. European Cells and Materials 24: 278–91.

Klein-Nulend J, Bakker AD, Bacabac RG, et al. 2013. Mechanosensation and transduction in osteocytes. Bone 54: 182–90.

Kramer I, Halleux C, Keller H, et al. 2010. Osteocyte Wnt/β-catenin signaling is required for normal bone homeostasis. Molecular and cellular biology 30: 3071–85.

Krishnan V, Bryant HU, and MacDougald OA. 2006. Regulation of bone mass by Wnt signaling. J Clin Invest 116: 1202–9.

Kujoth GC, Hiona A, Pugh TD, et al. 2005. Mitochondrial DNA mutations, oxidative stress, and apoptosis in mammalian aging. Science (80-) 309: 481–4.

Kuznetsova A, Brockhoff PB, and Christensen RH. 2017. lmerTest package: tests in linear mixed effects models. Journal of statistical software 82: 1–26.

Lenth RV. 2022. emmeans: Estimated Marginal Means, aka Least-Squares Means.

Lerebours C, Weinkamer R, Roschger A, and Buenzli PR. 2020. Mineral density differences between femoral cortical bone and trabecular bone are not explained by turnover rate alone. Bone reports 13: 100731.

Levchuk A. 2015. In Silico Investigation of Bone Adaptation in Health and Disease. ETH Zurich Research Collection.

Levchuk A, Zwahlen A, Weigt C, et al. 2014. The Clinical Biomechanics Award 2012—presented by the European Society of Biomechanics: large scale simulations of trabecular bone adaptation to loading and treatment. Clinical biomechanics 29: 355–62.

Li X, Ominsky MS, Niu Q-T, et al. 2008. Targeted deletion of the sclerostin gene in mice results in increased bone formation and bone strength. Journal of Bone and Mineral Research 23: 860–9.

Li X, Zhang Y, Kang H, et al. 2005. Sclerostin binds to LRP5/6 and antagonizes canonical Wnt signaling. Journal of Biological Chemistry 280: 19883–7.

Lin C, Jiang X, Dai Z, et al. 2009. Sclerostin mediates bone response to mechanical unloading through antagonizing Wnt/β-catenin signaling. Journal of bone and mineral research 24: 1651–61.

Lodish H, Berk A, Zipursky L, et al. 1999. Identification and Purification of Cell-Surface Receptors. Molecular Cell Biology, WH Freeman, New York,.

Martin M, Sansalone V, Cooper DM, et al. 2019. Mechanobiological osteocyte feedback drives mechanostat regulation of bone in a multiscale computational model. Biomechanics and modeling in mechanobiology 18: 1475–96.

Nakahama K. 2010. Cellular communications in bone homeostasis and repair. Cellular and molecular life sciences 67: 4001–9.

Nakashima T, Hayashi M, Fukunaga T, et al. 2011. Evidence for osteocyte regulation of bone homeostasis through RANKL expression. Nature medicine 17: 1231–4.

Nelson CA, Warren JT, Wang MW-H, et al. 2012. RANKL employs distinct binding modes to engage RANK and the osteoprotegerin decoy receptor. Structure 20: 1971–82.

Pastrama M-I, Scheiner S, Pivonka P, and Hellmich C. 2018. A mathematical multiscale model of bone remodeling, accounting for pore space-specific mechanosensation. Bone 107: 208–21.

Paul GR. 2020. Individualised multiscale mechanoregulation of fracture healing in mice. ETH Zurich Research Collection.

Pearson OM and Lieberman DE. 2004. The aging of Wolff’s “law”: ontogeny and responses to mechanical loading in cortical bone. American journal of physical anthropology 125: 63–99.

Perier-Metz C, Cipitria A, Hutmacher DW, et al. 2022. An in silico model predicts the impact of scaffold design in large bone defect regeneration. Acta Biomaterialia.

Perier-Metz C, Duda GN, and Checa S. 2020. Mechano-biological computer model of scaffold-supported bone regeneration: effect of bone graft and scaffold structure on large bone defect tissue patterning. Frontiers in Bioengineering and Biotechnology 8: 1245.

Pistoia W, Rietbergen B Van, Lochmüller E-M, et al. 2002. Estimation of distal radius failure load with micro-finite element analysis models based on three-dimensional peripheral quantitative computed tomography images. Bone 30: 842–8.

Raggatt LJ and Partridge NC. 2010. Cellular and molecular mechanisms of bone remodelling. Journal of Biological Chemistry: jbc–R109.

Rauner M, Jähn K, Hemmatian H, et al. 2020. Cardiovascular Calcification and Bone Mineralization. In: Cellular Contributors to Bone Homeostasis. Springer.

Robling AG, Niziolek PJ, Baldridge LA, et al. 2008. Mechanical stimulation of bone in vivo reduces osteocyte expression of Sost/sclerostin. Journal of Biological Chemistry 283: 5866–75.

Rodan GA. 1998. Bone homeostasis. Proceedings of the National Academy of Sciences 95: 13361–2.

Santos A, Bakker AD, and Klein-Nulend J. 2009. The role of osteocytes in bone mechanotransduction. Osteoporosis international 20: 1027–31.

Satopää V, Albrecht J, Irwin D, and Raghavan B. 2011. Finding a” kneedle” in a haystack: Detecting knee points in system behavior. In: 2011 31st international conference on distributed computing systems workshops.

Scheuren AC. 2020. Longitudinal assessment of frailty and osteosarcopenia in an in vivo model of premature aging.

Scheuren AC, D’Hulst G, Kuhn GA, et al. 2020a. Hallmarks of frailty and osteosarcopenia in prematurely aged PolgA (D257A/D257A) mice. J Cachexia Sarcopenia Muscle 11: 1121–40.

Scheuren AC, Kuhn GA, and Müller R. 2020b. Effects of long-term in vivo micro-CT imaging on hallmarks of osteopenia and frailty in aging mice. PLoS One 15: e0239534.

Schulte FA, Ruffoni D, Lambers FM, et al. 2013a. Local mechanical stimuli regulate bone formation and resorption in mice at the tissue level. PLoS One 8: e62172.

Schulte FA, Zwahlen A, Lambers FM, et al. 2013b. Strain-adaptive in silico modeling of bone adaptation - A computer simulation validated by in vivo micro-computed tomography data. Bone 52: 485–92.

Sculley D. 2010. Web-scale k-means clustering. In: Proceedings of the 19th international conference on World wide web.

Shahnazari M, Dwyer D, Chu V, et al. 2012. Bone turnover markers in peripheral blood and marrow plasma reflect trabecular bone loss but not endocortical expansion in aging mice. Bone 50: 628–37.

Strang G. 1968. On the construction and comparison of difference schemes. SIAM journal on numerical analysis 5: 506–17.

Tang Y, Wu X, Lei W, et al. 2009. TGF-β1-induced migration of bone mesenchymal stem cells couples bone resorption with formation. Nature medicine 15: 757–65.

Tol AF van, Schemenz V, Wagermaier W, et al. 2020. The mechanoresponse of bone is closely related to the osteocyte lacunocanalicular network architecture. Proceedings of the National Academy of Sciences 117: 32251–9.

Tourolle D. 2019. A Micro-scale Multiphysics Framework for Fracture Healing and Bone Remodelling. ETH Zurich Research Collection.

Tourolle DC, Dempster DW, Ledoux C, et al. 2021. Ten-Year Simulation of the Effects of Denosumab on Bone Remodeling in Human Biopsies. JBMR plus 5: e10494.

Trifunovic A, Wredenberg A, Falkenberg M, et al. 2004. Premature ageing in mice expressing defective mitochondrial DNA polymerase. Nature 429: 417–23.

Trüssel AJ. 2015. Spatial mapping and high throughput microfluidic gene expression analysis of osteocytes in mechanically controlled bone remodeling. ETH Zurich Research Collection.

Viceconti M, Juárez MA, Curreli C, et al. 2019. Credibility of in silico trial technologies—A theoretical framing. IEEE journal of biomedical and health informatics 24: 4–13.

Warren JT, Zou W, Decker CE, et al. 2015. Correlating RANK ligand/RANK binding kinetics with osteoclast formation and function. Journal of cellular biochemistry 116: 2476–83.

Webster DJ, Morley PL, Lenthe GH van, and Müller R. 2008. A novel in vivo mouse model for mechanically stimulated bone adaptation-a combined experimental and computational validation study. Comput Methods Biomech Biomed Engin 11: 435–41.

Weivoda MM, Ruan M, Pederson L, et al. 2016. Osteoclast TGF-β receptor signaling induces Wnt1 secretion and couples bone resorption to bone formation. Journal of Bone and Mineral Research 31: 76–85.

Winkler DG, Sutherland MK, Geoghegan JC, et al. 2003. Osteocyte control of bone formation via sclerostin, a novel BMP antagonist. The EMBO journal 22: 6267–76.

Xiong J, Piemontese M, Onal M, et al. 2015. Osteocytes, not osteoblasts or lining cells, are the main source of the RANKL required for osteoclast formation in remodeling bone. PLoS One 10: e0138189.

You L, Temiyasathit S, Lee P, et al. 2008. Osteocytes as mechanosensors in the inhibition of bone resorption due to mechanical loading. Bone 42: 172–9.

