## Supplementary Material for "Trabecular bone remodeling in the ageing mouse: a micro-multiphysics agent-based *in silico* model using single-cell mechanomics"

**Abbreviations:**

| Abbreviation | Description |
|---|---|
| AMGCL | Algebraic multigrid solver |
| ANOVA | Analysis of variance |
| BFR | Bone formation rate |
| BRR | Bone resorption rate |
| BS/BV | Specific bone surface |
| BS/TV | Bone surface density |
| BV/TV | Trabecular bone volume fraction |
| CSCS | Swiss National Supercomputing Center |
| CV6 | Sixth caudal vertebra |
| dt | Timestep |
| EFF | Effective strain |
| ES | Eroded surface |
| FQR | Formed Quiescent and Resorbed |
| GCC | Greatest Connected Component |
| HA | Hydroxyapatite |
| HSC | Hematopoietic stem cell |
| IPL | Image Processing Language |
| IVD | Intervertebral disc |
| LRP5/6 | Lipoprotein receptor-related protein 5/6 |
| MAR | Mineral apposition rate |
| $\text{mech}_{\text{thres}}$ and $mech_{thres}$ | Value of the mechanical signal leading to a production level corresponding to half of the maximum production capacity of a cell |
| MRR | Mineral resorption rate |
| micro-CT | Micro-computed tomography |
| micro-FE | Micro-finite element analysis |
| micro-MPA | Micro-multiphysics agent-based |
| MPI | Message Passing Interface |
| MS | Mineralizing surface |
| MSC | Mesenchymal stem cells |
| n | Number of samples |
| n.s | Not significant |
| Norm. BS/BV | Normalized specific bone surface |
| Norm. BS/TV | Normalized bone surface density |
| Norm. BV/TV | Normalized trabecular bone volume fraction |
| Norm. Tb.N | Normalized trabecular number |
| Norm. Tb.Sp | Normalized trabecular spacing |
| ParOSol | Parallel Octree Solver |
| PolgA | DNA polymerase subunit gamma A |
| pre-Oc | Pre-osteoclast |

| | |
|---|---|
| pre-Ot | Pre-osteocyte |
| OPG | Osteoprotegerin |
| Ob | Osteoblast |
| Oc | Osteoclast |
| ODE | Ordinary differential equation |
| OpenMP | Open multiprocessing |
| Ot | Osteocytes |
| RANK | Receptor activator of nuclear factor $\kappa$β |
| RANKL | Receptor activator of nuclear factor $\kappa$β ligand |
| RDD | Reaction, Diffusion, Decay |
| Scl | Sclerostin |
| Tb.N | Trabecular number |
| Tb.Sp | Trabecular spacing |
| Tb.Th | Trabecular thickness |
| TGF-β1 | Transforming growth factor beta 1 |
| β | Maximum production value by a single cell |
| ε | Gaussian-dilated effective strain |

### 1 Supplementary Data

The time step of the cells follows the execution order of the different actions from the original version of the model. They are executed in the following order:

Move Cells: MSCs, HSCs; Cell Proliferation; Cell Death; Preosteocyte to Osteocyte; Osteoclast surface behavior; Preosteoclast surface behavior; HSC to preosteoclast; Cell Productions; Osteoblast Movement; Osteoblast surface behavior; Matrix Mineralization; MSC to osteoblast; Lining cell-osteoblast differentiation.

#### 1.1 Single-cell mechanomics

The single-cell mechanomics is defined as the synthesis of cytokines in the voxel where the cell is present or the amount of bone or collagen to be removed by a single osteoclast. Compared to the original implementation of the cell productions, we employ the Hill function to model the single-cell mechanomics response as a function of the local mechanical signal. The following definitions were applied to osteoblasts, osteoclasts and osteocytes. The anabolic and anti-catabolic mechanomics is defined as the following increasing function f(ε):

$$f(\varepsilon) = \frac{\beta}{1 + \left(\frac{mech_{thres}}{\varepsilon}\right)^{n}}$$

Where β is the maximum production value by a single cell, $mech_{thres}$ : Value of the mechanical signal corresponding to β/2, ε is the gaussian-dilated effective strain, n is the Hill coefficient. Analogously, the anti-anabolic and catabolic mechanomics is defined as the following mechanomics decreasing function g(ε):

$$g(\varepsilon) = \beta \cdot \left(1 - \frac{1}{1 + \left(\frac{mech_{thres}}{\varepsilon}\right)^n}\right)$$

#### 1.2 Subdivision of 3D lattice

The subdivision of the 3D lattice was obtained using the following pseudocode. Here, we optimized the number of divisions along each axis to achieve the lowest total area of voxels that have to be exchanged between subdomains.

Example of pseudocode:

```
INT MPI_size ← 32
INT shapex ← 200
INT shapey ← 200
INT shapez ←300
INT min_area ← shapex*shapey*shapez*MPI_size
FOR cx = 1 to MPI_size
   FOR cy = 1 to MPI_size
      FOR cz = 1 to MPI_size
         INT  product ← cx*cy*cz
         IF product = MPI_size THEN
            INT area ← shapey*shapez* (cx-1)+ shapex*shapez* (cy-1) + shapey*shapex* (cz-1)
            IF area < min_area THEN
               INT grid ← [cx, cy, cz]
               INT min_area ← area
            END IF
         END IF
      END FOR
   END FOR
END FOR
```

Here, the *MPI_size*, *shapex*, *shapey*, *shapez* are the number of MPI ranks (from 8 Nodes · 4 MPI tasks per node) and the dimension of the lattice along the x, y and z axis. The *grid* variable contains the number of subdivisions along the x, y and z axes, respectively. For example, a lattice of 200x200x300 voxels would have 2, 4 and 4 subdivisions along the x, y, and z axes, respectively. Therefore, each subdomain will have a size of 100x50x75 voxels, meaning 375 thousand voxels in each subdomain.

##### 1.3 Statistical models

Three linear mixed models (t1, t2, t3) were tested using the lmerTest package with these equations:

t1<-lmerTest::lmer(x ~ Data * time + (1|mouse_no), data=data)

t2<-lmerTest::lmer(x ~ Data * time + (Data | mouse_no), data=data)

t3<-lmerTest::lmer(x ~ time + (Data | mouse_no), data=data)

Where "x" is a static or dynamic parameter. "Data" is either *in vivo* measurement or *in silico* simulation when comparing the simulation to the *in vivo* data. Also, "Data" is either the group of simulations with high, medium or low production of OPG, RANKL and Scl when comparing the three production levels of a given cytokine. "mouse_no" is the ID of a mouse. "Time" is the measurement time, from 0 to 4 weeks, for the static parameters. Also, "Time" is the baseline time point, either 0 or 2 weeks, for the dynamic parameters. t2 model was eventually chosen based on model section criteria (*Bayesian information criteria)* for performing the statistical tests.

#### 2 Supplementary Tables

Table S1: Reference configuration of the parameters of the in silico model.

| Parameter | Description | Value |
|---|---|---|
| $D$ | Diffusivity of the cytokines | $2.0 \times 10^{-8}\ cm^2/s$ |
| $\lambda^{\mathrm{TGF-\beta1}}$ | Decay of TGF-β1 | $0\ 1/s$ |
| $\lambda$ | Decay of all the other cytokines | $4.0 \times 10^{-6}\ 1/s$ |
| $k^{f}_{LRP5/6,Scl}$ | Forward binding coefficient for LRP5/6-Scl | $0.95 \times\ 10^{-1}\ 1/(zmol\ s)$ |
| $k^{r}_{LRP5/6,Scl}$ | Backward binding coefficient for LRP5/6-Scl | $1.0 \times\ 10^{-4}\ 1/s$ |

| | | |
|---|---|---|
| $k^{f}_{\mathrm{TGF-\beta1\_rec,TGF-\beta1}}$ | Forward binding coefficient for TGF-β1_rec- TGF-β1 | $2.0 \times 10^{-2}\ 1/(zmol\ s)$ |
| $k^{r}_{\mathrm{TGF-\beta1\_rec,TGF-\beta1}}$ | Backward binding coefficient for TGF-β1_rec- TGF-β1 | $1.0 \times 10^{-3}\ 1/s$ |
| $k^{f}_{RANK,RANKL}$ | Forward binding coefficient for RANK-RANKL | $4.0 \times 10^{-1}\ 1/(zmol\ s)$ |
| $k^{r}_{RANK,RANKL}$ | Backward binding coefficient for RANK-RANKL | $1.0 \times 10^{-3}\ 1/s$ |
| $k^{f}_{OPG,RANKL}$ | Forward binding coefficient for OPG-RANKL | $1.0 \times 10^{-1}\ 1/(zmol\ s)$ |
| $k^{r}_{OPG,RANKL}$ | Backward binding coefficient for OPG-RANKL | $8.0 \times 10^{-4}\ 1/s$ |
| $n_{Oc}$ | Hill coefficient osteoclast | 5 - |
| $n_{Ob}$ | Hill coefficient osteoblast ad lining cell | 7 - |
| $n_{Ot}$ | Hill coefficient Osteocyte | 5 - |
| $\omega_{Ob}$ | Osteoblast random movement probability | 0.2 - |
| $\omega_{Pre.Oc}$ | Preosteoclast random movement probability | 45% - |
| $\omega^{RANKL}_{Pre.Oc}$ | Preosteoclast biased movement to neighbor with highest RANKL | 25% - |
| $\tau^{LRP5/6}$ | LRP5/6 bound receptor threshold: above this value MSCs (if they can) and | 70% |

| | | |
|---|---|---|
| | osteoblasts differentiate into lining cells | |
| $\tau^{RANK}$ | RANK bound receptor threshold: above this values HSCs (if they can) differentiate into preosteoclasts | 50% |
| $\tau^{\mathrm{TGF-\beta1}}$ | TGF-β1 bound receptor threshold: above this values MSCs and osteoblasts (if they can) have a proliferation scale increased by the factor $s_{TGF\_beta1}$ | 12% |
| $\sigma_{Ob}$ | Osteoblast polarization | 0.65 - |
| $\sigma_{Oc}$ | Osteoclast polarization | 0.65 - |
| $Clus_{Ob}$ | Osteoblast cluster size | 3 - |
| $Clus_{Oc}$ | Osteoclast cluster size | 3 - |
| $r^{mineral}$ | Mineralization rate | 2.5 $1/(day)$ |
| $\beta_{Ot}^{Scl}$ | Max osteocyte anti-anabolic production rate of Scl | $1.0 \times 10^{-3} zmol\ /20\ mins$ |
| $\beta_{Ot}^{RANKL}$ | Max osteocyte catabolic production rate of RANKL | $0.25 \times 10^{-2} zmol\ /20\ mins$ |
| $\beta_{Ot}^{OPG}$ | Max osteocyte anti-catabolic production rate of OPG | $0.3 \times 10^{-2} zmol\ /20\ mins$ |
| $P_{MSC}$ | MSC proliferation rate | 0.04 $1/day$ |

| | | |
|---|---|---|
| $P_{HSC}$ | HSC proliferation rate | $0.08\ 1/day$ |
| $P_{Ob}$ | Osteoblast proliferation rate | $0.01\ 1/day$ |
| $P_{PreOc}$ | Preosteoclast proliferation rate | $0.08\ 1/day$ |
| $A_{Ot}$ | Osteocyte apoptosis rate | $6.8 \times 10^{-5}\ 1/day$ |
| $A_{Ob}$ | Osteoblast apoptosis rate | $0.01\ 1/day$ |
| $A_{Oc}$ | Osteoclast apoptosis rate | $0.5 \times 10^{-2}\ 1/day$ |
| $\beta_{Oc}^{mineral,osteoid}$ | Max osteoclast catabolic resorption rate of mineral and osteoid | $0.5\ zmol/day$ |
| $\beta_{Ob}^{osteoid}$ | Max osteoblast anabolic synthesis rate of osteoid | $0.1\ zmol/day$ |
| $\beta_{Ob,Li.Ce}^{RANKL}$ | Max osteoblast and lining cell catabolic production rate of RANKL | $0.4 \times 10^{-3} zmol\ /20\ mins$ |
| $\beta_{Ob,Li.Ce}^{OPG}$ | Max osteoblast and lining cell anabolic production rate of OPG | $0.6 \times 10^{-2} zmol\ /20\ mins$ |
| $s^{\mathrm{TGF-\beta 1}}$ | TGF-β1 proliferation scaling factor | 2 |
| $\mathrm{dt}_{\mathrm{cells\text{-}RDD}}$ | Timestep for the update of the positions and state of the cells as well as for the reaction-diffusion-decay of the cytokines | 40 minutes |

| | | |
|---|---|---|
| $dt_{micro\text{-}FE}$ | Timestep for the update of the mechanical signal and of the TGF-β1 diffusion matrix | 8 hours |

Table S2: Initial values of receptor activator of nuclear factor κB ligand (RANKL), osteoprotegerin (OPG), RANKL-OPG, sclerostin (Scl) and transforming growth factor beta 1 (TGF-β1) concentrations used in the simulation.

| Cytokine | Value |
|---|---|
| RANKL | 102 pg/ml (top and bottom third of the lattice) |
| | 0.051 pg/ml (middle third of the lattice) |
| OPG | 1100 pg/ml (top and bottom third of the lattice) |
| | 11 pg/ml (middle third of the lattice) |
| RANKL-OPG | 1166 pg/ml |
| Scl | 72 pg/ml (top and bottom third of the lattice) |
| | 0.72 pg/ml (middle third of the lattice) |
| TGF-β1 | 7 pg/ml (in marrow) |
| | 21 pg/ml (in bone) |

Table S1: Low, basal and high single-cell production values of osteoprotegerin (OPG) by the osteocytes.

| | **OPG** |
|---|---|

| | |
|---|---|
| Low value | $0.258 \times 10^{-2} zmol\ /20\ mins$ |
| Basal value (reference for comparisons) | $0.3 \times 10^{-2} zmol\ /20\ mins$ |
| High value | $0.342 \times 10^{-2} zmol\ /20\ mins$ |

Table S2: Low, basal and high single-cell production values of receptor activator of nuclear factor kB ligand (RANKL) by the osteocytes.

| | **RANKL** |
|---|---|
| Low value | $0.202 \times 10^{-3} zmol\ /20\ mins$ |
| Basal value (reference for comparisons) | $0.25 \times 10^{-2} zmol\ /20\ mins$ |
| High value | $0.298 \times 10^{-3} zmol\ /20\ mins$ |

Table S3: Low, basal and high single-cell production values of sclerostin (Scl) by the osteocytes.

| | **Scl** |
|---|---|
| Low value | $0.787 \times 10^{-3} zmol\ /20\ mins$ |
| Basal value (reference for comparisons) | $1.0 \times 10^{-3} zmol\ /20\ mins$ |
| High value | $1.213 \times 10^{-3} zmol\ /20\ mins$ |

Table S6: Full report of two-way ANOVA analysis of the static and dynamic bone morphometry between *in vivo* and *in silico* data.

| | Group | Time | Group x Time |
|---|---|---|---|
| BV/TV | n.s. | p<0.01 | p<0.01 |
| BS/BV | p<0.01 | p<0.0001 | p<0.0001 |
| BS/TV | p<0.05 | p<0.0001 | p<0.0001 |
| Tb.Th | n.s. | p<0.01 | n.s. |
| Tb.Sp | p<0.05 | p<0.0001 | p<0.0001 |
| Tb.N | p<0.001 | p<0.0001 | p<0.0001 |
| BFR | n.s. | p<0.05 | p<0.01 |
| BRR | n.s. | n.s. | n.s. |
| MAR | p<0.01 | p<0.001 | p<0.001 |
| MRR | p<0.001 | p<0.01 | p<0.05 |
| MS | p<0.0001 | n.s. | p<0.05 |
| ES | p<0.0001 | n.s. | n.s. |

Table S7: Full report of two-way ANOVA and post-hoc analyses of the static and dynamic bone morphometry between the different OPG levels.

| | ANOVA | | | Post-hoc analysis (Group comparison) | | |
|---|---|---|---|---|---|---|
| | Group | Time | Group x Time | High-low | High-basal | Basal-low |
| NormBVTV | p<0.01 | p<0.0001 | p<0.0001 | p<0.01 | n.s. | p<0.01 |

| | | | | | | |
|---|---|---|---|---|---|---|
| NormBSBV | p<0.001 | p<0.0001 | p<0.05 | p<0.01 | n.s. | p<0.01 |
| NormBSTV | n.s. | p<0.0001 | n.s. | n.s. | n.s. | n.s. |
| NormTbTh | p<0.001 | p<0.01 | n.s. | p<0.01 | n.s. | p<0.05 |
| NormTbSp | n.s. | p<0.0001 | n.s. | n.s. | n.s. | n.s. |
| NormTbN | n.s. | p<0.0001 | n.s. | n.s. | n.s. | n.s. |
| BFR | n.s. | p<0.0001 | n.s. | n.s. | n.s. | n.s. |
| BRR | p<0.0001 | p<0.0001 | p<0.05 | p<0.01 | p<0.05 | p<0.05 |
| MAR | n.s. | p<0.0001 | n.s. | n.s. | n.s. | n.s. |
| MRR | n.s. | p<0.0001 | n.s. | n.s. | n.s. | n.s. |
| MS | p<0.05 | p<0.01 | n.s. | n.s. | p<0.01 | p<0.05 |
| ES | p<0.01 | n.s. | p<0.05 | p<0.01 | p<0.05 | p<0.05 |

Table S8: Full report of two-way ANOVA and post-hoc analyses of the static and dynamic bone morphometry between the different RANKL levels.

| | ANOVA | | | Post-hoc analysis (Group comparison) | | |
|---|---|---|---|---|---|---|
| | Group | Time | Group x Time | High-low | High-basal | Basal-low |
| NormBVTV | p<0.0001 | p<0.01 | p<0.0001 | p<0.001 | p<0.001 | p<0.01 |
| NormBSBV | p<0.0001 | p<0.0001 | p<0.0001 | p<0.001 | p<0.01 | p<0.05 |
| NormBSTV | n.s. | p<0.0001 | n.s. | n.s. | n.s. | n.s. |
| NormTbTh | p<0.0001 | p<0.01 | p<0.0001 | p<0.001 | p<0.05 | n.s. |

| | | | | | | |
|---|---|---|---|---|---|---|
| NormTbSp | n.s. | p<0.0001 | n.s. | n.s. | n.s. | n.s. |
| NormTbN | p<0.01 | p<0.0001 | p<0.0001 | p<0.05 | p<0.01 | p<0.05 |
| BFR | p<0.01 | p<0.0001 | p<0.01 | p<0.05 | n.s. | p<0.05 |
| BRR | p<0.0001 | p<0.01 | p<0.001 | p<0.001 | p<0.05 | p<0.001 |
| MAR | p<0.05 | p<0.0001 | n.s. | n.s. | n.s. | n.s. |
| MRR | n.s. | p<0.0001 | n.s. | n.s. | n.s. | n.s. |
| MS | p<0.001 | p<0.0001 | p<0.01 | p<0.001 | p<0.05 | p<0.01 |
| ES | p<0.001 | p<0.05 | p<0.0001 | p<0.001 | p<0.001 | p<0.01 |

Table S9: Full report of two-way ANOVA and post-hoc analyses of the static and dynamic bone morphometry between the different Scl levels.

| | ANOVA | | | Post-hoc analysis (Group comparison) | | |
|---|---|---|---|---|---|---|
| | Group | Time | Group x Time | High-low | High-basal | Basal-low |
| NormBVTV | p<0.001 | p<0.0001 | n.s. | p<0.01 | p<0.01 | n.s. |
| NormBSBV | p<0.0001 | p<0.0001 | n.s. | p<0.01 | p<0.05 | n.s. |
| NormBSTV | p<0.05 | p<0.0001 | n.s. | n.s. | p<0.05 | n.s. |
| NormTbTh | p<0.01 | p<0.001 | n.s. | p<0.05 | p<0.05 | n.s. |
| NormTbSp | p<0.05 | p<0.0001 | n.s. | n.s. | p<0.05 | n.s. |
| NormTbN | n.s. | p<0.0001 | n.s. | n.s. | n.s. | n.s. |

| | | | | | | |
|---|---|---|---|---|---|---|
| BFR | p<0.01 | p<0.0001 | n.s. | n.s. | p<0.05 | n.s. |
| BRR | n.s. | p<0.0001 | n.s. | n.s. | n.s. | n.s. |
| MAR | n.s. | p<0.0001 | n.s. | n.s. | n.s. | n.s. |
| MRR | n.s. | p<0.0001 | n.s. | n.s. | n.s. | n.s. |
| MS | p<0.001 | p<0.0001 | p<0.0001 | p<0.05 | p<0.01 | p<0.05 |
| ES | n.s. | n.s. | n.s. | n.s. | n.s. | n.s. |
